# Krüppel-like factors KLF5 & KLF8 emerge as potential master transcriptional regulators of Alzheimer’s disease, as revealed on cell fate regulomes in human brain organoids

**DOI:** 10.64898/2026.07.23.740401

**Authors:** Antoine Aubert, Anne-Claire Comby, Aude Bramoulle, Maria Grazia Mendoza-Ferri, Peggy Azzolin, Huan Li, Alice Moussy, Sudeshna Das, Bruno Maria Colombo, Marco Antonio Mendoza-Parra

## Abstract

In addition to the well described beta-amyloid plaques accumulation and tau hyper- phosphorylation, Alzheimer’s disease (AD) is accompanied by major changes in gene expression. Herein, we aimed at revealing master transcription factors (TFs) responsible for gene expression changes during AD progression. For this, we have used human brain organoids (BORGs) harbouring AD-related genetic mutations (APP-Swedish, PSEN1- M146V), which were traced over multiple time-points by bulk and spatially-resolved transcriptomics. By reconstructing gene regulatory networks (GRNs) and applying a machine learning framework that classifies transcription factors (TFs) based on predicted regulatory cascade dynamics, we identified a core set of 64 AD specific master TFs. These factors were subsequently validated by their over expression in transcriptomes from human patients.

Strikingly, both KLF5 and KLF8 showed promoter enrichment across more than half of the AD specific transcription factors, strongly suggesting that they occupy key regulatory positions within the AD regulome. This functional relevance was further emphasized by spatially resolved transcriptomics, which demonstrated that their regional over expression mirrors the spatial distribution of AD markers in human patient samples.

Finally, we have demonstrated that this AD-specific regulome is controlled by the aberrant CREB3L2-ATF4 heterodimer previously described as being induced by the beta-amyloid plaques deposition. Overall, these findings reconstitute the regulome behind the progression of AD and highlights key TFs as potential druggable targets for the treatment of the disease.

## INTRODUCTION

Alzheimer’s disease (AD) is the most common form of dementia affecting ∼ 40 million people worldwide (Tahami Monfared et al. 2022). In addition to the beta-amyloid plaques and tau hyper-phosphorylation aggregates accumulation in the brain, AD is accompanied by major changes in gene expression, arguing for the necessity of pairing these events to enhance our understanding of the disease. Importantly, these types of studies require to count with model systems allowing to interrogate for the progression of the disease, instead of interrogating the end-point status like in the case of post-mortem samples, but in addition to conserve the human setting.

In the last years, the use of in-vitro 3-dimensional cell cultures presenting self-organized properties – known as cerebral or brain organoids (BORGs) – is revolutionizing the study of human nervous tissue development. Human induced pluripotent stem cells (hIPSCs) can be differentiated in nervous tissue, composed by a variety of neuronal subtypes, but also macroglia related cells. BORGs have been used for modeling neurodevelopmental events, but also to replicate key hallmarks of AD pathophysiology, boasting amyloid plaques and neurofibrillary tangles when using hIPSCs harboring early-onset AD-related familial mutations (e.g. APP-Swedish (Gonzalez et al. 2018; Raja et al. 2016) ; PSEN1 mutations (Arber et al. 2021; Vanova et al. 2023)), or late-onset AD genetic risk factors (e.g. APOE4 (Zhao et al. 2020); Bin1 (Saha et al. 2024)).

Considering that BORGs allow to perform longitudinal studies from the hIPSC state till the formation of complex cortical layer structures, in this study we aimed at exploiting this capacity to trace the emergence of gene regulatory programs defining the AD state setup. For this we took advantage of human iPSCs harboring AD-genetic mutations and analyze them by bulk transcriptomics across 120 days of BORG differentiation. These data were used for retrieving master Transcription factors (TFs) specifically enriched on AD BORGs thanks to the use of a machine learning strategy, which were then validated both in human AD patients’ sample transcriptomes as well as in spatially-resolved transcriptomic assays performed on 5- month BORGs harboring the APP-Swedish mutation.

This integrative effort revealed a subset of 64 master TFs, specifically up-regulated in AD. Furthermore, the Krüppel-like factors KLF5 & KLF8 emerged as key regulators of the AD-specific TF co-regulatory network. Finally, the reconstructed AD-specific regulome has been integrated into a model focused on the induction of a deregulated AD-specific gene regulatory program driven by the formation of an aberrant CREB3L2-ATF4 heterodimer in the context of Beta amyloid plaques exposure.

## RESULTS

### Human Brain organoids harbouring familial Alzheimer’s disease genetic mutations present divergent cell differentiation progression

With the aim of studying the influence of familial Alzheimer’s disease (AD) genetic mutations in nervous tissue progression we took advantage of a panel of isogenic APP and PSEN1mutant hIPSCs previously generated by CRISPR/Cas9 genome editing (Kwart et al. 2019). Specifically, we have used hIPSCs harbouring the APP Swedish mutation (*APP-swe*), the Methyonine-Valine mutation in the amino acid 146 retrieved within the PSEN1 gene (*PSEN1-M146V*), as well as an isogenic *APP-swe/PSEN1-M146V* “double mutant” line. These hIPSCs followed a cortical brain organoid (BORG) culture protocol, based on the use of a triple-inhibitors action (Dual SMAD, TGFB and WNT inhibition ( Roseborck et al (Rosebrock et al. 2022))) leading to an homogeneous and reproducible 3-dimensional nervous tissue differentiation (**Supplementary Figure S1**). BORGs issued from AD-related genetic background as well as their isogenic WT counterpart were characterized over-time by immune-staining measurements, bulk transcriptomes as well as spatially-resolved transcriptomic assays at the late stages of differentiation (**Figure 1A**).

**Figure 1.**
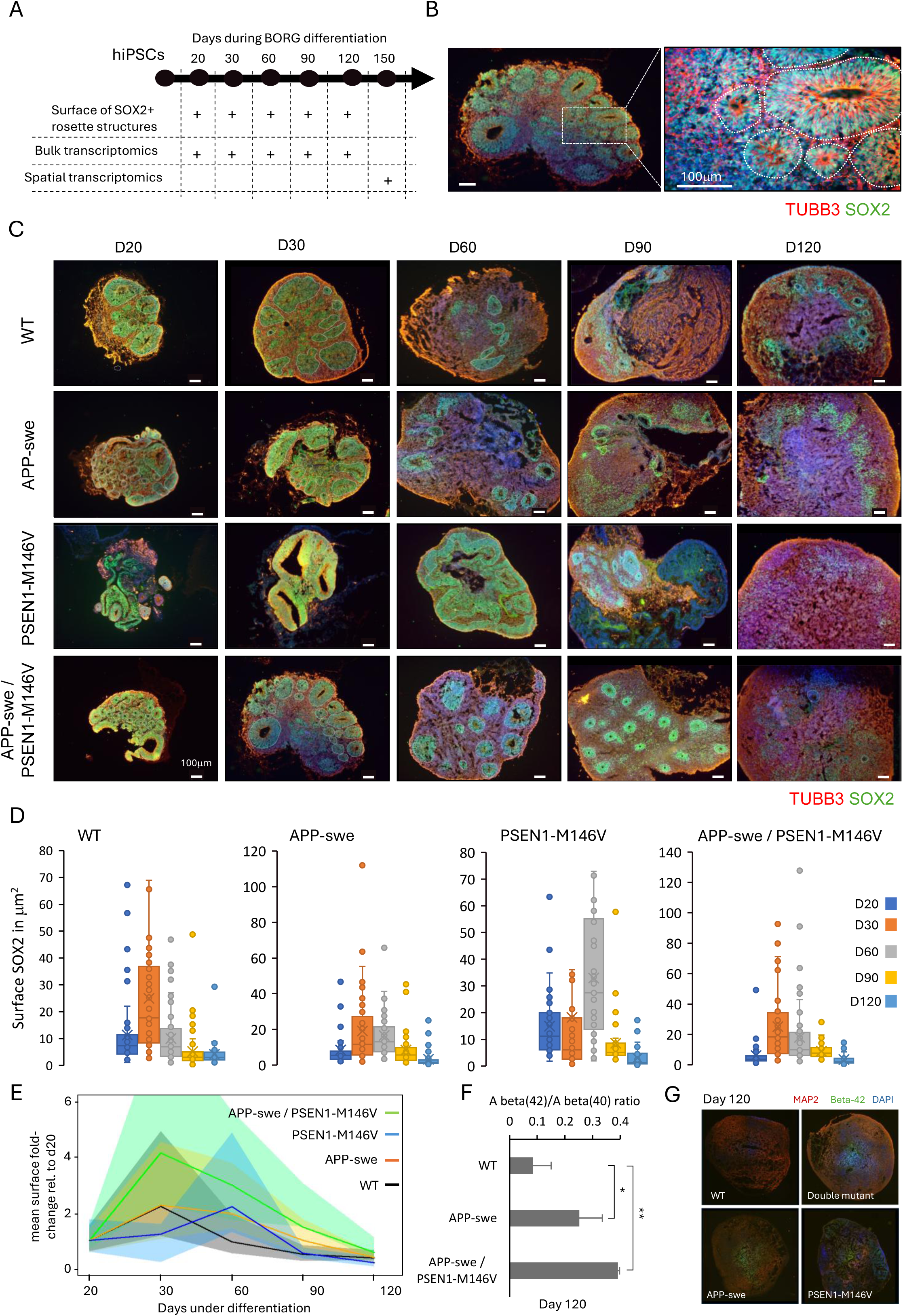
Human induced pluripotent stem cells (hIPSCs) harboring familial Alzheimer’s disease (AD) genetic mutations present a divergent neural progenitor cells progression during Brain organoids (BORGs) differentiation (A) Scheme illustrating samples collection during BORGs differentiation and the applied analytical readout. (B) Immunofluorescence micrograph of a BORG section revealing the presence of the neuronal marker TUBB3 (red) and the pluripotent neuronal stem cell marker SOX2 (green). Inset provides a magnified view where the SOX2-positive rosette radial arrangements were demarcated as a measure of the surface occupied by neural progenitor cells in the tissue. (C) Representative immunofluorescence micrographs for the markers TUBB3 and SOX2 assessed on tissue sections issued from WT and AD-related BORGs and collected at various stages during differentiation. As indicated in (B), the observed rosette-like structures were demarcated per section and used to evaluate the surface occupied by neural progenitor cells at different stages during BORGs progression. (D) Boxplot display of the surface of SOX2- positive rosette-like structures assessed at various time-points during BORGs differentiation. (E) Average SOX2-positive rosette-like structures surface normalized relative to the measurement assessed at day 20. The displayed shaded confidence intervals are delimited by the Q75 and Q25 quantiles. (F) Amyloid Beta 42/40 ratios determined on BORGs cultured over 120 days issued from the APP-swe or the APP-swe/PSEN1-M146V double mutant line in comparison with their isogenic control WT line. Welch’s ANOVA test P-value: 0.003255 *p < 0.02, **p < 0.01 via adjusted t-test. (G) Representative immunofluorescence micrographs of BORG of 120 days revealing the presence of amyloid-beta aggregates (Beta- 42 in Green). Mature neurons are revealed by MAP2 (red). Displayed analyses are issued from two independent kinetic cultures in which at least three brain organoids per time-point are processed.

In the early phases of BORG differentiation, neurogenesis is characterized by the emergence of polarized neural rosette-like structures composed of neural progenitor cells (NPCs), as revealed by the presence of the Sex determining region Y-related HMG-box gene 2 (SOX2) marker (**Supplementary Figure S2** and **Figure 1**). Considering that NPCs give rise to all neural cell types, it is expected that the number and the size of rosette structures per BORG during differentiation provide a readout of their progression. By analyzing several SOX2- positive immune-stained sections (**Figure 1B**), we have confirmed a temporal gain of the rosette surface in WT BORGs during the first 30 days of differentiation (**Figure1C&D**). After this time-point a rapid decrease of the surface of the SOX2-positive rosette structures has been observed, which coincides with the emergence of differentiated cells like GFAP-positive astrocytes (**Supplementary Figure S3**).

Rosette surface analysis performed on BORGs harboring the PSEN1-M146V genetic mutation revealed a major gain after 60 days culture, while the analysis of BORGs harboring either the APP-swe mutation or the combination of both APP-swe and PSEN1-M146V revealed a peak after 30 days of differentiation but a slow decrease in size on the following time-points (**Figure 1C&D**).

The comparison of the rosette kinetics assessed in all lines revealed a similar BORG development progression on WT and both APP-swe and the double-mutant line during the first 30 days, but followed by a slow decrease for the AD lines. In contrary, the PSEN1- M146V line presented a delayed gain on size for the rosette structures, but followed by a rapid decrease (**Figure 1E**). These rosette surface kinetic discrepancies relative to the isogenic WT counterpart coincides with the delayed presence of GFAP-positive cells in the AD-mutant lines (**Supplementary Figure S3**). Despite these observations, BORGs harboring the AD-related mutations do present a significant gain on Amyloid Beta 42 after 120 days of differentiation (**Figure 1F&G**).

To have a more in depth-view of the molecular changes taking place during BORG development progression, we have generated bulk transcriptomic readouts from samples collected at different time-points during the first 120 days of differentiation. The differential gene expression response per time-point relative to readouts assessed from hIPSCs was stratified on gene co-expression paths. As illustrated in **Figure 2A**, a total of 15 gene co-expression paths were identified for the WT BORGs, from which 5 of them present significant differentially gene co-expression responses (|LFC|>3). A Gene Ontology (GO) analysis performed on the significant differentially gene co-expressed paths revealed that the early upregulated Path 1 (P.1: 231 genes up-regulated from day 20 onwards) is enriched for terms associated to neuronal differentiation and notably neural stem cells, while Path 2 (P.2: 308 genes; upregulated from day 30), Path3 (P.3: 590 genes; upregulated from day 90) or Path5 (P.5: 205 genes; bifurcated from the upregulation of path1 at day 30) are preferentially enriched on specialized cell types like astrocytes (P.1,P.2, P3), GABAergic or Glutamatergic neurons (P.3), as well as oligodendrocyte precursors (P.5) (**Figure 2B**).

**Figure 2.**
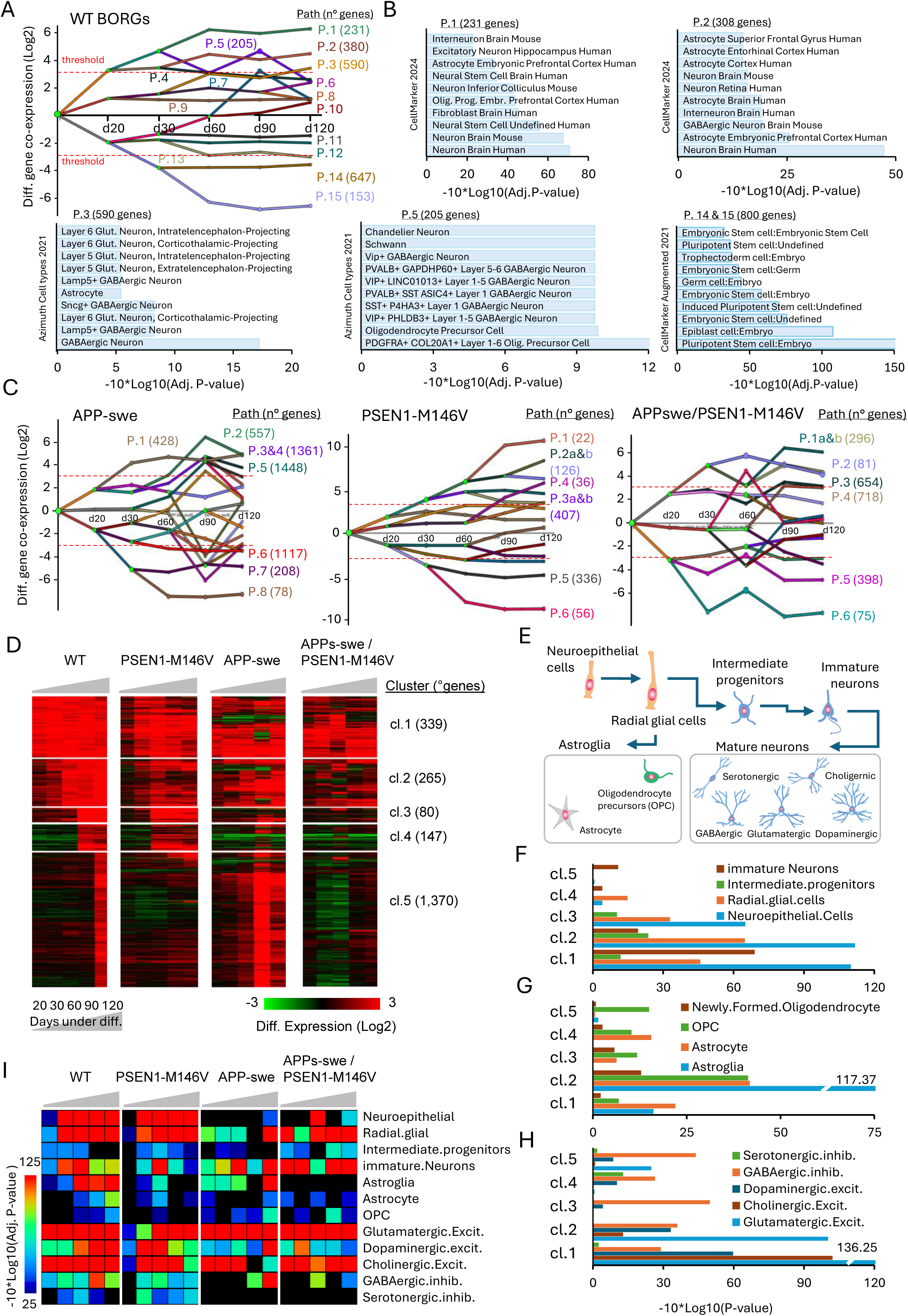
Discrepancies on cell fate progression during BORGs differentiation issued from hIPSCs harboring AD-related mutations. **(A)** Stratification of the temporal transcriptome profiling during WT BORGs differentiation in 15 gene co-expression paths, accompanied by relevant bifurcation points (green circles). For Paths by-passing the indicated fold-change threshold the number of genes per path are indicated. Dashed red line delineates the LFC threshold of +/-3 used for defining significant differentially expressed paths. **(B)** Gene Ontology terms enrichment analysis performed on the significant paths. **(C)** Same as (A) but performed on BORGs harboring AD-related genetic mutations. **(D)** Up- regulated genes associated to WT BORGs differentiation were classified on 5 clusters based on their temporal enrichment; and their expression per cluster on BORGs harboring the AD- related mutations were displayed. **(E)** Scheme displaying the neural cell lineage expected during BORGs progression. **(F-H)** Cell fate probabilities associated to each gene clusters displayed in (D). **(I)** Cell fate probabilities assessed in a temporal manner for BORGs harboring the AD-related mutations as well as for the isogenic WT counterpart.

In contrast to WT, gene co-expression maps assessed on AD-related BORGs revealed a delayed response for the earliest up-regulated paths (up-regulated response (LFC>3) from day 30 onwards; **Figure 2C**). A direct comparison between the GO terms enrichment confidences assessed on gene co-expression paths retrieved on WT and AD-related BORGs confirmed the presence of neuronal differentiation signatures within the early up-regulated gene co-expression paths but with a weaker confidence for BORGs harboring the PSEN1-M146V mutation (**Supplementary Figure S4**). Furthermore, the later up-regulated paths in AD- BORGs presented GO terms enrichment associated to specialized cell types like in WT conditions, but presenting slight differences. Notably, the GO enrichment confidences associated to Oligodendrocyte precursor cells (OPC) were rather low or inexistent for the AD-related BORGs, arguing for a delayed differentiation progression relative to their WT counterparts (**Supplementary Figure S4**). This is further supported by the fact that a direct differential expression analysis between AD-related BORGs relative to the WT counterpart at day 120 revealed an important number of deregulated genes (1,466 genes up-regulated in PSEN1-M146V; 1,036 in APP-swe; and 768 genes in the case of the double-mutant line relative to WT counterpart; **Supplementary Figure S5A**). Furthermore, a rather low number of common genes between the different AD lines were retrieved (47 common up-regulated and 85 common down-regulated genes; **Supplementary Figure S5B**), arguing for a different differentiation state after 120 days between WT BORGs and those harboring AD-related mutations.

To better decorticate BORG development progression, we have stratified all up-regulated genes based on their temporal induction. In WT BORGs five different gene clusters were identified, composed by genes induced as soon as 20 days of treatment (cl.1: 339 genes), genes induced between 30-60 days of treatment (cl.2: 265 genes), genes induced after 90 days of treatment (cl.3: 80 genes; cl.4: 147 genes) and genes being up-regulated only after 120 days of culture (cl.5: 1,370 genes) (**Figure 2D**). Gene expression response on AD-related BORGs based on WT clusters revealed the loss of up-regulated signatures associated to the later time- point (day 120; cluster 5) for BORGs harboring the PSEN1-M146V or the APPs-swe/PSEN1- M146V mutations. Furthermore, BORGs harboring the APP-swe mutation presented an earlier induction for genes retrieved in cluster 5 (**Figure 2D**). To understand the consequences of such differences, we have used a cell-type specific gene set enrichment analysis targeting neural precursor cell fate during WT BORG development progression (**Figure 2E** **& Supplementary Table S1**). This analysis revealed the strong enrichment on neuroepithelial cells in cluster1 and cluster 2, and the gain of enrichment confidence for radial glial cells and intermediate progenitors in cluster 2 (**Figure 2F**). Furthermore, cluster 2 were strongly enriched for signatures associated to Astroglia (absent in later stage clusters), further accompanied by the presence of signatures associated to Astrocyte and OPCs (**Figure 2G**). Finally, cluster 1&2 presented signatures associated to Glutamatergic and cholinergic neurons, while GABAergic neurons appeared enriched on all clusters (**Figure 2H**).

By applying the cell-type specific gene set enrichment analysis per time-point instead of per cluster, we can also confirm that signatures associated to neuroepithelial and radial glial cells are retrieved between day 30 and day 120 on WT BORGs and those harboring the PSEN1- M146V mutation (**Figure 2I**). This if further supported by the expression signatures observed for the radial gial cell gene markers FABP7, HES1, PAX6 or SLC1A3 (**Supplementary Figure S6**). Astroglia signatures in WT are observed from day 60, and Astrocyte or OPC signatures are retrieved at day 120. Importantly, glial cell signatures are rather absent or delayed (APP-swe) in AD-related BORGs (**Figure 2I**). Indeed, the astrocyte markers GFAP and S100B are seen significantly induced in WT BORGs from day 60, while detected in APP- swe BORGs at day 120 (**Supplementary Figure S6 & Supplementary Figure S3)**.

Finally, immature neuron signatures are transitory enriched on WT BORGs between day 30- day 60 (while lasting at lower levels till day 120) while delayed (day 60) for PSEN1-M146V and APP-swe. Specialized glutamatergic or cholinergic neuron markers are either retrieved enriched from day 20 onwards (neurons) in all BORG types (confirmed by the expression of the Glutamatergic neuron markers SLC17A6 and GRIN2B; **Supplementary Figure S6**), while Dopaminergic neuron markers are enriched from day 60 in WT BORGs, mainly in day 20-day 30 in PSEN1-M146V, and strongly delayed in APPswe or the double mutant line. Furthermore, GABAergic markers are observed from day 90 on WT BORGs but either absent or delayed in AD-related BORGs (**Figure 2I**).

Overall, while WT and AD-related BORGs present neural precursor cell types including neuroepithelial and radial glial cells during the first 120 days of differentiation, a delay in the emergence of mature cell types like Astrocytes or specialized neurons is observed in AD BORGs.

### A network of master transcription factors is specifically retrieved on BORGs harbouring Alzheimer’s disease-related mutations

Stratification of differential gene expression readouts collected during BORGs differentiation in co-expressed paths revealed the presence of bifurcation decision points (**Figure 2A**). Gene co-expression bifurcation events are meant to be the consequence of transcription factors (TF) action (Ernst et al. 2007; Schulz et al. 2012). Previously we have developed TETRAMER (Cholley et al. 2018; Mendoza-Parra et al. 2016), a computational strategy able to combine differential gene expression readouts with transcription factor (TF)-target gene (TGs) relationships assessed from public databases to (i) build gene regulatory networks (GRNs) conserving TF-TG relationships that are coherent with the differential gene expression response of the system of interest; (ii) rank TFs based on their capacity to control down- stream TGs in a direct or indirect manner. This machine-driven strategy scrutinizes the capacity of each TF within the reconstructed GRN to propagate a transcription regulation signalling downstream within the connectivity and then compare its performance with randomized GRNs to evaluate its significance. As consequence, the ranking strategy is able to identify most influent TFs, called “master TFs” within the system (**Figure 3A**).

**Figure 3.**
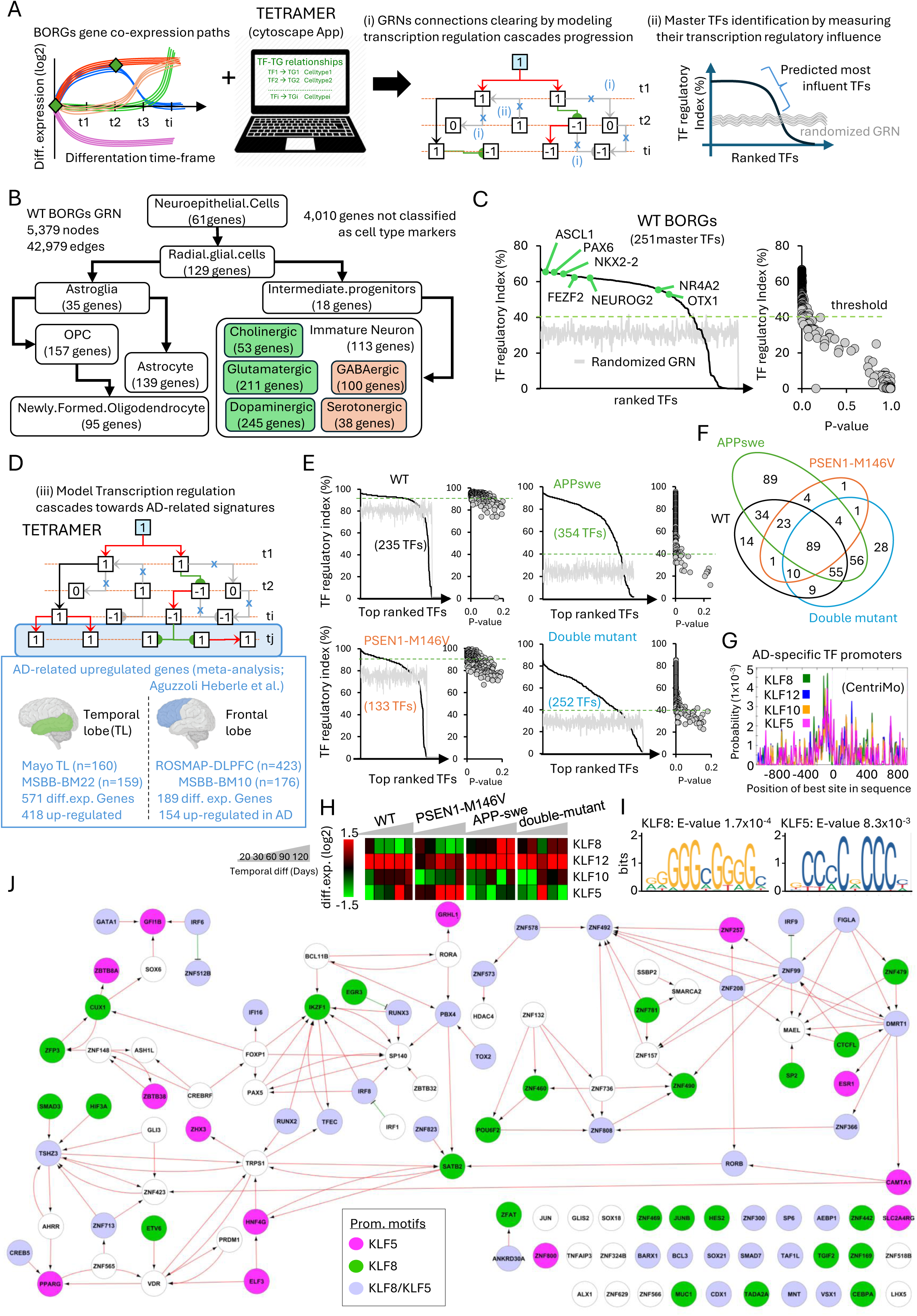
A network of master transcription factors is specifically deregulated on BORGs harboring Alzheimer’s disease-related mutations. **(A)** Scheme illustrating the integration of temporal gene co-expression paths assessed on BORGs with a collection of Transcription factors-Target gene (TF-TG) relationships issued from public databases. This integrative effort is performed within TETRAMER, a Cytoscape Application allowing to reconstruct a gene regulatory network (GRN) from the above information (Cholley et al. 2018). TETRAMER propagates transcription regulation cascades from each of the TFs retrieved within the GRN, and detects incoherent TF-TG relationships based on the differential gene expression status (1,-1, or 0 for induced, repressed or non-differentially expressed) at each analyzed time-points (t1,t2…ti) and the type of transcription regulation (gray arrow: activation; green line: repression). Hence, (i) edges presenting an incoherence between the gene expression status between the interconnected nodes (genes); or (ii) nodes interconnections that are not rising from the in silico activated TF are removed from the GRN. Furthermore, TETRAMER classifies all TFs based on the fraction of downstream TGs that are controlled within the system and evaluates the confidence of this classification by performing the same analysis on randomized GRNs. **(B)** Representation of the GRN reconstituted by TETRAMER from the temporal transcriptomes assessed on wild-type BORGs. In addition to indicate the number of total nodes (genes) and edges (relationships) composing the GRN, the presence of gene markers associated to various neural cell types retrieved during BORGs progression are displayed. **(C)** Left panel: Ranked TFs based on the fraction of downstream target genes under their control (TF regulatory index in percent). Gray line corresponds to the average TF regulatory index issued from 20 randomized GRNs. Right panel: P-value confidence assessed from the comparison between the assessed TF regulatory index issued from the reconstructed GRN and that observed in a randomized context. **(D)** Modeling transcription regulation cascades leading to the activation of genes known to be up-regulated in AD patients’ samples processed as part of the meta-analysis performed by Heberle et al. (Aguzzoli Heberle et al. 2025). For Temporal lobe (TL) samples: Mayo Clinic samples (160 datasets); Mount Sinai Brain Bank cohort (MSBB) Brodmann area in use: 22 (MSBB-BM22; 159 datasets). For Frontal lobe samples: ROSMAP DLPFC (dorsolateral prefrontal cortex; 423 datasets); Mount Sinai Brain Bank cohort (MSBB) Brodmann area in use: 10 (MSBB-BM10; 176 datasets). For this analysis the 418 up-regulated genes in AD in temporal lobe as well as the 154 up-regulated genes in frontal lobe samples were merged and used as final targets within the transcription regulation cascades modeling performed by TETRAMER. **(E)** Ranked TFs based on their capacity to activate the upregulated genes issued from (D). Green dashed line corresponds to the threshold in use for selecting significant master TFs. Notice that this threshold is defined by the randomized GRN analysis **(F)** Venn-diagram displaying the common and specific master transcription factors revealed in (E). **(G; I)** Motif analysis performed on promoter regions of the AD-specific TFs found in (F), revealing the enrichment of the transcription factors KLF8, KLF12, KLF10 and KLF5. CentriMo confidence for KLF8: E = 1.7 × 10^-4^; KLF5: E = 8.3 × 10^-3^. **(H)** temporal differential expression of KLF factors in BORGs harboring either the indicated AD-genetic mutations or in the isogenic wild-type control background. Timepoints under study: 20,30,60,90 and 120 days. **(J)** Gene co-regulatory network displaying the transcription regulation relationship between the AD-specific TFs found in (F). Red arrowed edges: activation; green edges: repression. Nodes are colored based on the presence of either a KLF5, KLF8 or both biding site motifs within their promoters.

By applying this strategy on the temporal transcriptomics data assessed on WT BORGs, we obtained a GRN composed of 5,379 nodes (genes) and >42,900 connections (edges), including ∼1,000 gene markers associated to the neural precursor cell fate, i.e. from neuroepithelial, until glial cells or specialized neurons (**Figure 3B**).

Ranking TFs within WT BORG progression based on their regulatory capacity revealed the presence of 251 master TFs (yield>40%; i.e. controlling >40% of the downstream target genes), among them ASCL1 (a well-known neural differentiation factor essential for cell type diversification (reviewed in Lundie-Brown et al. 2025)), PAX6 (known as human neuroectoderm cell fate determinant factor (Zhang et al. 2010)), NKX2-2 (homeodomain protein that plays a main role in oligodendrocyte differentiation (Li et al. 2009), and motor neurons cell fate (Jarrar et al. 2015)), FEZF2 (a transcriptional repressor essential for the differentiation of subcortical projection neurons (Zuccotti et al. 2014)), NEUROG2 (known as a potent neuronal differentiation factor (Hulme et al. 2022)), NR4A2 (widely expressed in the central nervous system and known to play crucial roles in various processes like dopaminergic neuronal differentiation, or synaptic plasticity (Duan et al. 2026)), or OTX1 (a key transcription factor regulating cortical neurogenesis (Huang et al. 2018)) (**Figure 3C**).

A similar analysis performed on differential gene expression readouts assessed on AD-related BORGs revealed 215 master TFs in APP-swe, 225 TFs in PSEN1-M146V and 239 TFs in BORGs harboring both the APP-swe and PSEN1-M146V mutations (**Supplementary Figure S7A&B)**. Among all these identified master players, a total of 86 common TFs were identified among all BORG samples, while 160 TFs are specific to the AD BORGs (**Supplementary Figure S7C**). Importantly, while the common TFs were shown to be involved in controlling neuronal differentiation, the AD-specific TFs corresponded to developmental transcription factors known to be repressed in neural precursors or specialized nervous tissue; arguing for their aberrant activation (**Supplementary Figure S7D&E**).

Considering that the studied mutations are known AD genetic drivers, we aimed at evaluating the presence of gene programs associated to this disease within the studied BORGs. For this, we have taken advantage of an AD-related up-regulated gene expression collection established from human patients’ samples issued from the temporal (418 genes) or the frontal lobe (154 genes) (Aguzzoli Heberle et al. 2025). TETRAMER has been used for interrogating the presence of these up-regulated genes within BORGs and notably to identify the master TFs that would be able to activate such AD patients-related gene panel (**Figure 3D**). This analysis revealed 235 TFs in WT BORGs as able to control the up-regulation of the AD patients-related gene panel and notably with a rather low confidence (P-value<0.2). In contrast, a total of 354 highly confident master TFs were found in the case of the APP-swe mutation (P-vaule<0.05), 133 TFs for PSEN1-M146V (P-value<0.2), and 252 TFs for BORGs harbouring both mutations (P-value<0.12) (**Figure 3E**). When comparing all identified master TFs, we found a common panel of 89 TFs, and a group of 183 TFs specifically associated to the AD-related BORGs (**Figure 3F**). A motif analysis performed on the promoter region of the 183 AD-specific TFs revealed a high enrichment of four motifs associated to the Krüppel-like factor family, namely KLF8, (known to be highly expressed in the mouse neurons (Dobrivojević et al. 2015)), KLF12 (known to be expressed in various tissues including brain, kidney, liver or lung (Pan et al. 2023)), KLF10 (shown to be expressed in various regions of the developing and adult brain in the mouse and known to regulate the emergence of both astrocyte and oligodendrocyte lineages (Garduño-Tamayo et al. 2025)), and KLF5 (recently shown to play a main role in maintaining the neural precursor cell (NPC) populations, regulate their proliferation and neuronal differentiation (Fuchigami et al. 2026)). (**Figure 3G**). Gene expression profiling assessed for these four TFs relative to the hIPS state revealed the over-expression of KLF12 within all lines; KLF10 appeared non-differentially expressed in WT BORGs, while downregulated in AD BORGs. KLF5 appeared transitory induced in WT BORGs (day 90) while strongly over-expressed over-time in the PSEN1-M146V mutant BORG (**Figure 3H** and **Supplementary Figure S8**). Finally, KLF8 appeared specifically induced in BORGs harbouring the APP-swe mutation (**Figure 3H** and **Supplementary Figure S8**).

Based on the AD-specific over-expression behaviour observed for KLF8 and KLF5, supported by their high confidence associated to the binding motif detection (**Figure 3I**), we have reconstructed a KLF8/KLF5 regulome by considering the most significant AD-specific master TFs. As illustrated in **Figure 3J**, KLF8 / KLF5 motifs were found in 75 of the 110 AD-specific master TFs (selected by AD-specific up-regulation from the 183 AD-specific master TFs) composing the illustrated co-regulatory network, strongly arguing for their important role in driving the AD-related programs retrieved within BORGs. Notably, KLF8/KLF5 motifs were associated to TFs like GATA1, previously shown to mediate transcriptional regulation of the gamma-secretase activating protein (GSAP)(Chu, Wisniewski, et Praticò 2016); IRF6 (Interferon regulatory factor 6), known to be expressed in the brain, but in addition its up-regulation has been associated to neuronal apoptosis promotion (Lin et al. 2016); the cyclic-AMP response binding protein 5 (CREB5), known to play a key role in shifting amyloid processing by repressing the metalloproteinase ADAM10 and activating the beta-site cleaving enzyme 1 (BACE1) (Zeng et al. 2025); or RUNX2, a TF previously found to be over-expressed in AD, leading to a reduction of neuronal differentiation in favour of astrocytes (Nakatsu et al. 2023).

Similarly, KLF8 motifs were found in promoter regions associated to genes like the hypoxia-inducible transcription factor HIF3A, previously shown to mediate an aberrant oxidative stress response in AD (Xue et al. 2026); or the zinc finger transcription factor ZNF460, recently proposed as a pivotal regulator of the neuronal stress response in AD (Gong et al. 2025).

KLF5 motifs were found in genes like ELF3, known to be required for neural tissue development in zebrafish (Sarmah et al. 2022), PPARG, described recently as a key regulator associated to ferroptosis and neuroinflammation in AD (Tang et al. 2026), or CAMTA1, a calcium-responsive TF known to be strongly expressed in the brain, thought to regulate neuroprotective genes under stress; and presenting epigenetic dysregulation in both brain and blood before onset of sever amyloidosis in mice models (Okhovat et al. 2025).

### AD-specific master transcription factors revealed in BORGs are over-expressed in AD-patients’ samples issued from various brain regions

Since brain organoids are per definition in-vitro generated tissue reconstituting neurodevelopmental material, the pertinence of the AD-specific regulome revealed in BORGs requires to be confirmed in real AD patients’ samples. For this, we have interrogated publicly available transcriptomics data issued from AD cohorts (Alzheimer’s disease DataLENS (Noori et al. 2024)). Specifically, we have interrogated datasets issued from the Mayo clinic Brain Bank (MayoBB TCX: Temporal cortex; CBE: Cerebellum); The Mount Sinai Brain Bank (MSBB IFG: Inferior Frontal Gyrus; STG: Superior Temporal Gyrus; FP: Frontal pole; PHG: Parahippocampal Gyrus), as well as those issued from the ROSMAP consortium (PFC: prefrontal cortex).

Gene set enrichment analysis (GSEA) performed between the master TFs found specifically up-regulated in AD-BORGs (**Figure 3J**) and the AD-upregulated signatures retrieved in the aforementioned patients’ sample collections revealed that 64 of the 110 TFs presented significant enrichment levels within at least one of the interrogated databases (**Figure 4A**). Indeed, most of the 64 enriched TFs present a significant up-regulated enrichment in AD patients’ samples issued from either temporal cortex (TCX) or cerebellum (CBE) samples retrieved within the Mayo Brain Bank collection (**Figure 4B**), while a fraction of them is significantly upregulated in either the Mount Sinai Brain Bank samples collection or within the ROSMAP cohort (**Figure 4B**).

**Figure 4.**
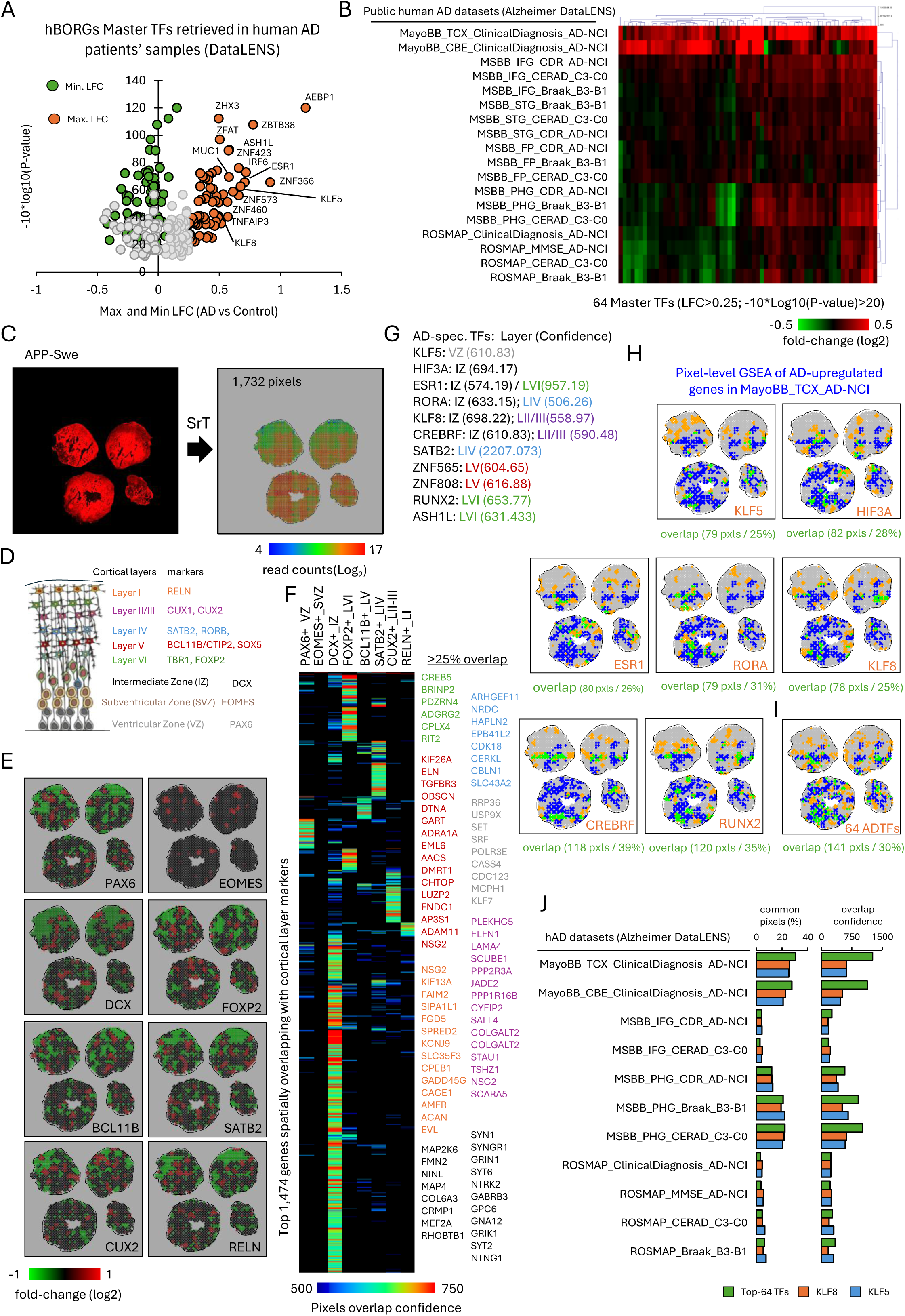
AD-specific master transcription factors revealed in BORGs are over- expressed in AD-patients’ samples issued from various brain regions. Master TFs found specifically over-expressed in hBORGs harboring AD-specific genetic mutations were interrogated for their differential expression in human AD patients’ samples and ranked by Gene set enrichment analysis (GSEA) leading to the assessment of an adjusted P-value confidence. GSEA analysis has been performed in 18 different datasets issued from various patients’ cohorts retrieved within the Alzheimer’s DataLENS database (Noori et al. 2024). **(A)** Maximum (orange) and minimum (green) significant log2 fold-change (LFC) expression levels associated to the interrogated TFs as retrieved across the various AD patients’ sample datasets displayed relative to their adjusted P-value confidence levels issued from the GSEA enrichment analysis. Gray cercles correspond to non-significantly deregulated TFs. A total of 64 Significant TFs were identified, being defined as LFC >0.25 and a -10*Log10(P-value)>20 **(B)** Heatmap displaying the LFC associated to the 64 master TFs selected in (A) in the context of each of the interrogated human AD patients’ sample datasets. MayoBB: Mayo clinic brain bank; TCX: Temporal cortex; CBE: Cerebellum; NCI: No cognitive impairment control samples; MSBB: Mount Sinai brain bank; IFG: Inferior Frontal Gyrus; STG: Superior Temporal Gyrus; FP: Frontal Pole; PHG: Parahippocampal Gyrus; CDR: Clinical Dementia Rating; Braak: Braak Stage (score focused on tau neurofibrillary tangles) B3 (stage V/VI; neocortical) vs B1 (stage 0; normal, I/II (hippocampal); CERAD: CERAD stage (Neuropathological score focused on the frequency of Beta-amyloid neuritic plaques) C3 (Frequent/Definite AD) vs C0 (None / Not AD); MSE: Mini Mental State Examination; ROSMAP: Samples issued from the Religious Order Study and Memory and Aging Project. **(C)** Spatially-resolved transcriptomics (SrT) profiling assay performed on human brain organoids (hBORGs) of 5 months harboring the APP-swe mutation. Left panel: Tissue section corresponding to four hBORGs labelled by dCTP-Cy3 incorporation during cDNA production over in-house manufactured double-barcoded DNA arrays (Lozachmeur et al. 2023). Right panel: SrT profile composed by 1,732 pixels. Heatmap corresponds to the total read counts per pixel expressed in Log2 scale. **(D)** Schematic representation of the embryonic cerebral tissue proliferation from the ventricular zone (VZ) towards the formation of the various layers composing the human brain cortex. In addition, gene markers associated to each layer are indicated. Adapted from (Mukhtar et Taylor 2018). **(E)** Spatial gene expression of the various gene markers associated to the different human cortex layers as retrieved within the analyzed APP-swe BORGs. Heatmap correspond to the local differential gene expression relative to the average behavior over the whole surface under study. **(F)** Top spatially over-expressed genes overlapping with the various BORG regions associated to the indicated cortex layers. Heatmap corresponds to the hypergeometric probability associated to the overlap between the interrogated genes and the gene marker associated to the indicated cortex layer. Besides the heatmap, the indicated genes correspond to top ranked genes based on their hypergeometric overlapping probability and presenting >25% of overlap with the gene marker. Genes are colored in agreement with the cortical layer color code displayed in (D). **(G)** AD-specific TFs initially retrieved as part of the selected subset in (A)&(B) and presenting significant overlapping confidence with the various cortical layer regions within APP-swe hBORGs. **(H)** Pixel-level gene set enrichment analysis (GSEA) revealing AD-related regions across the APP-swe hBORGs (MayoBB TCX AD-NCI dataset) compared with the spatial regions presenting the over-expression of the TFs listed in (G). Green pixels: overlapping regions; Blue pixels: regions enriched on genes associated to the MayoBB TCX database; Orange pixels: regions presenting the over-expression of the indicated TF. At the bottom of each panel the number of overlapping pixels and the fraction (in percentage) relative to the total number of pixels per TF are displayed. **(I)** Same as (H) but displaying the comparison with regions occupied by at least one of the 64 AD-specific TFs. **(J)** Spatial overlapping between either the 64 AD-specific TFs, KLF8 or KLF5 and the spatial regions enriched (pixel level GSEA) on genes up- regulated in the indicated AD patients’ sample datasets. Left panel: fraction of common pixels (in percentage); Right panel: Hypergeometric overlap confidence.

Since such differences in TF expression within AD patients’ samples could be due to the brain regions under study, we thought that in the same manner, TF expression within BORGs should be evaluated by taking in consideration tissue complexity. For this reason, we have cryo-sectioned multiple BORGs collected after 5 months of differentiation and analyse them by using our in-house developed spatially-resolved transcriptomics (SrT) (Lozachmeur et al. 2023). As illustrated in **Figure 4C**, four BORGs harbouring the APP-swedish mutation were processed by SrT and gave rise to a digitized map composed by 1,732 pixels. Considering that the BORGs differentiation protocol in use is expected to generate cortical tissue, and that the comparison with public transcriptomics data issued from AD patients’ samples presented a strong enrichment correlation in temporal cortex material, we have first confirmed the up- regulation of gene markers associated to the various cortical layers (as reviewed by (Mukhtar et Taylor 2018)) (**Figure 4D**). Indeed, as illustrated in **Figure 4E**, markers associated to the ventricular zone (VZ) like PAX6, the subventricular zone (SVZ) like EOMES, or the intermediate zone like DCX were retrieved as upregulated within the BORG tissue sections. Furthermore, expression of markers associated to the layer VI (FOXP2), layer V (BCL11B/CTIP2), layer IV (SATB2), layer II/III (CUX2), and layer I (RELN) were detected across the BORG sections (**Figure 4E**). To further support this analysis, we have evaluated the spatial co-expression of other genes at the regions where the aforementioned cortex layer markers were observed. Indeed, a total of 1,474 genes were significantly spatially co-enriched with at least one of the aforementioned markers. Importantly, 166 genes were co-enriched with the PAX6+ regions, including among them various canonical or strongly VZ-associated genes, like RRP36 (ribosome biogenesis, progenitor-enriched), USP9X (RG polarity and adhesion regulator), SET (chromatin regulator active in proliferative zones), SRF (progenitor cytoskeletal regulation), POLR3E (polymerase III subunits, enriched in proliferative zones), CASS4 (adhesion/cytoskeletal signaling in progenitors), CDC123 (translation initiation factor enriched in dividing cells), MCPH1 (translation initiation factor enriched in dividing cells), or KLF7; another Krüppel-like transcription factor known to regulate neurogenesis and neuronal migration (Hong et al. 2023; Liu et al. 2025) (**Supplementary Table S2** & **Figure 4F**).

Similarly, 994 genes appeared significantly spatially over-expressed within the DCX+ region, including 42 canonical IZ markers, 23 IZ Neuronal signalling/Synaptic-priming genes (**Supplementary Table S2** & **Figure 4F**). Other 300 genes were spatially co-expressed with the FOXP2+LVI; 107 genes with the BCL11+LV; 263 genes with the SATB2+LIV; 180 genes with the CUX2+_LII/III; and 75 genes with the RELN+_LI (**Supplementary Table S2** & **Figure 4F**). In all cases, previously described markers for the aforementioned cortex layers were systematically retrieved spatially co-expressed within the SrT data (**Supplementary Table S2** & **Figure 4F**).

Among the various retrieved genes associated to the aforementioned cortex layer regions, we have identified 11 TFs presenting a high overlapping confidence which are in addition part of the 64 up-regulated genes within AD-specific BORGs as well as within the AD-patients’s cohort samples. Among them we found the Krüppel-like factor 5 (KLF5), known to play a main role in maintaining neuronal precursors by suppressing their differentiation, as well as to promote the differentiation of PAX6+ radial glia cells in the ventricular zone (VZ) to the EOMES+ subventricular zone (SVZ) (Fuchigami et al. 2026). Furthermore, the hypoxia-inducible transcription factor HIF3A, the estrogen receptor encoding gene ESR1, the RAR-related orphan receptor alpha (RORA), the Krüppel-like factor 8 (KLF8), the CREB3 regulatory factor (CREBRF), presented significant gene co-expression confidence enrichment levels within the intermediate zone (IZ) (**Figure 4G**). Finally, SATB2, the canonical marker of layer IV; the zinc finger TFs ZNF565 & ZNF808, presenting significant gene co-expression enrichment levels within the layer V; and the transcription factors RUNX2 and ASH1L; both being enriched preferentially in layer VI; were retrieved specifically up- regulated in AD-related BORG (**Figure 4G**).

With the aim of revealing the BORG regions that are concerned by AD-specific signatures, we have performed a pixel-level gene set enrichment analysis (GSEA) against the Alzheimer’s DataLENS collection. This analysis performed on APP-swe BORGs revealed up to 468 pixels from the 1,732 (27%) enriched in AD signatures retrieved up-regulated within the Mayo Brain Bank temporal cortex datasets (**Supplementary Figure S9**); while less than 8% of the pixels (87 from 1,021) composing a SrT dataset assessed from Wild-type control BORG samples were observed for the same public human AD samples datasets (**Supplementary Figure S10**). Furthermore, a direct comparison between the SrT readouts assessed on the APP-swe BORGs and those issued from its isogenic counterpart WT control samples revealed a total of 1,205 common up-regulated genes, for 2,069 specifically up- regulated in WT and other 2,963 specific to the APP-swe BORGs (**Supplementary Figure S11**). A GO enrichment analysis performed on the common group of genes revealed the enrichment for markers associated to specialized neurons (GABAergic neurons, pyramidal cells; Panglao DB), as well as to nervous tissue regions (Prefrontal cortex, spinal cord: ARCHS4 tissues DB). Furthermore, the GO enrichment analysis performed on the APP-swe specific genes revealed the presence of terms like “Alzheimer’s disease” (WikiPathway 2021 DB), as well as “APP processing in Alzhemier’s disease” (Elsevier Pathway collection DB), confirming the presence of AD markers over-expressed in the APP-swe BORGs (**Supplementary Figure S11D**).

By evaluating the degree of overlap between the spatial gene expression of the aforementioned 11 AD-specific TFs with that of the pixels associated to AD signatures across the APP-swe BORG sections, we have found that 25% of the KLF5 over-expressed pixels do match with those presenting an AD-signatures enrichment issued from the MayoBB TCX datasets (**Figure 4H**). Similarly, HIF3A presented an overlap of 28%; 26% in the case of ESR1; 31% for RORA; 25% for KLF8; 39% for CREBRF; and 35% for RUNX2 (**Figure 4H**). Importantly, when considering all spatial regions over-expressing at least one of the 64 TFs, we have found that 30% of them spatially overlapped with the AD signatures issued from the MayoBB TCX datasets (**Figure 4I**). By extending this analysis to other human AD patients’ sample datasets retrieved within the Alzheimer DataLENS we have found 27% common pixels with the Mayo Brain Bank Cerebellum (CBE) collection, and 20 to 22% overlap when comparing the 64 AD-specific TFs with the Mount Sinai Brain Bank (MSBB) PHG (Parahippocampal Gyrus) datasets (AD vs the control samples based on Braak stage (B3-B1) or CERAD (C3-C0) scoring respectively) (**Figure 4J**). Furthermore, this spatial correlation analysis extended to the KLF8 or the KLF5 TFs revealed similar overlapping results (**Figure 4J**).

### Integrating the AD-specific up-regulated regulome found in BORGs within a model whereby CREB3L2-ATF4 aberrant heterodimer promotes the setup of AD-linked gene expression changes

By using human brain organoids harbouring AD-related genetic mutations we have managed to identify an aberrant regulome composed by at least 64 TFs, which were further confirmed by interrogating publicly available human patients’ samples cohorts (**Figure 4**). Among these TFs, a subset of 11 factors appeared as central players, from which both KLF5 and KLF8 appeared as key co-regulatory factors, notably due the presence of their binding motifs associated to > 60% of the other TFs (**Figure 3** **&** **Figure 4G**). Despite these findings, we still miss to understand the mechanism that connects the over-expression of this master TFs to the presence of the genetic mutations linked to AD that are part of the studied BORG tissues.

Recently, Gouveia Roque and colleagues demonstrated that the exposure of neurons to oligomerized Amyloid Beta 42 induce the aberrant heterodimerization of the CREB3L2- ATF4 transcription factors. Furthermore they validated the presence of such aberrant heterodimer on AD brain patients’ samples (Gouveia Roque et al. 2023). To reveal the regulome that is controlled by this aberrant heterodimer, Gouveia Roque and colleagues engineered a chemically induced proximity strategy relying on recombinant TF constructs composed by dimerization domains (FRB or FKBP located at the carboxy terminus) as well as the presence of epitope tagging moieties (HA or V5) (**Figure 5A**). By integrating these constructs in HEK cells in the context of CREB3L2 and ATF4, they have managed to reconstitute either both homodimer situations as well as the aberrant CREB3L2-ATF4 and analyzed by chromatin immunoprecipitation by pulling either in the HA or V5 epitopes (**Figure 5A**).

**Figure 5.**
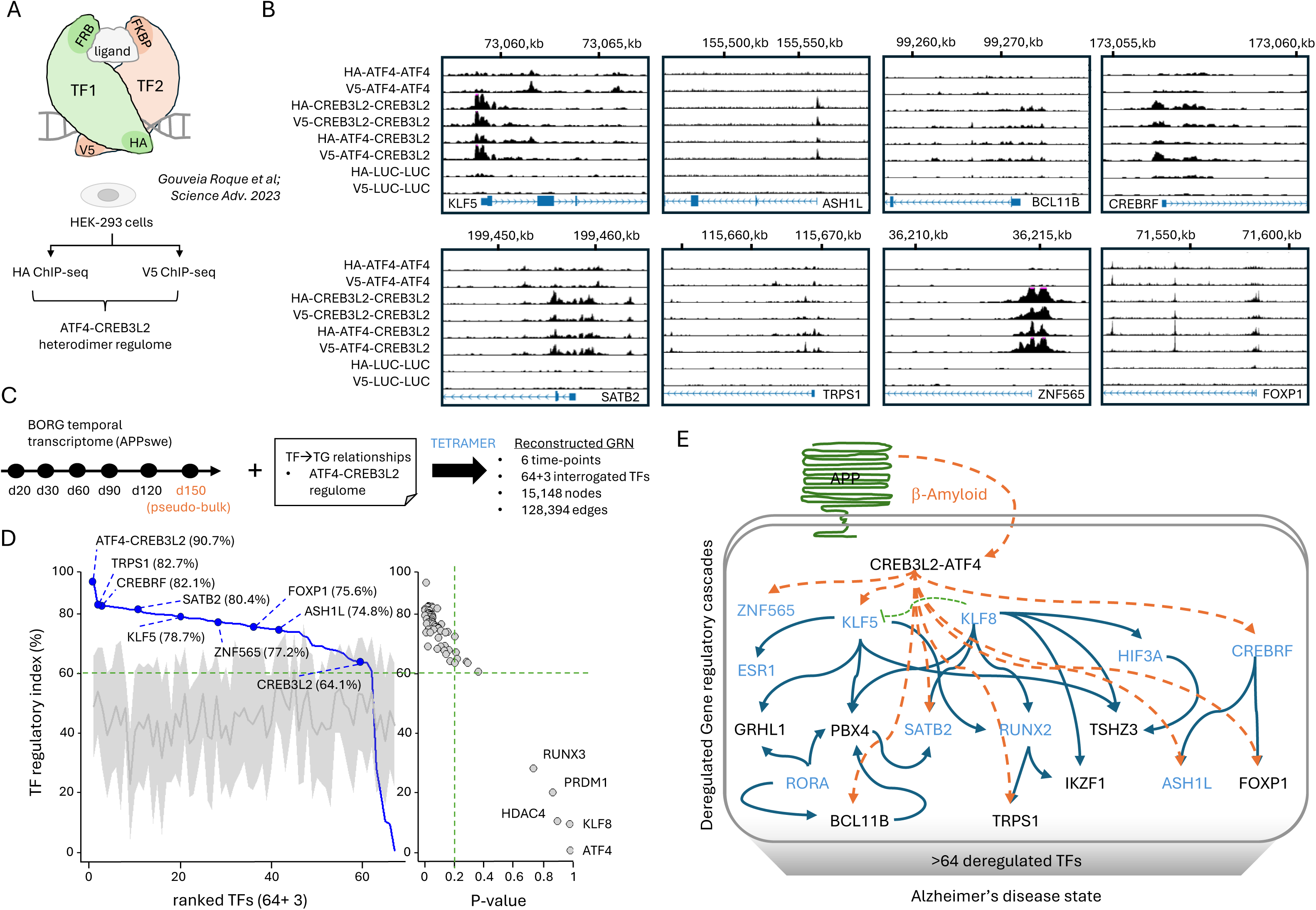
Integrating the AD-specific up-regulated regulome found in human BORGs within a model whereby CREB3L2-ATF4 aberrant heterodimer promotes the setup of AD-linked gene expression changes. **(A)** Scheme illustrating the “ChIPmera” strategy used by Gouveia Roque et al. for revealing the regulome associated to the aberrant CREB3L2- ATF4 heterodimer promoted by Beta-amyloid pathology. Top panel: Recombinant TF architecture based on the use of specific dimerization domains (FRB or FKBP) retrieved at the carboxy terminus, as well as a N-terminal epitope moiety (HA or V5) for efficient pull-down. Bottom panel: Human embryonic kidney (HEK) cells transfected with either homodimer or heterodimer constructs for the recombinant CREB3L2 and ATF4 were used for HA or V5 chromatin immunoprecipitation combined with massive parallel sequencing (ChIP-seq). As consequence, these ChIP-seq profiling maps are used for revealing the ATF4-CREB3L2 regulome. **(B)** CREB3L2-ATF4 heterodimer binding patterns observed in the promoter of representative TFs found as AD-specific in this study. The illustrated binding patterns are issued from the re-analysis of the ChIP-sequencing profiles performed by Gouveia Roque et al. with their chemically induced proximity strategy allowing to promote the formation of the indicated TF dimers (ChIPmera) (Gouveia Roque et al. s. d.). **(C)** Strategy for interrogating the performance of the ATF4-CREB3L2 heterodimer to control the panel of 64 master TFs associated to AD revealed in this study. Brain organoid (BORG) transcriptomes issued from days 20 till day 120 and complemented with a pseudo bulk transcriptome issued from the spatially-resolved transcriptome assessed at day 150 were integrated with the ATF4-CREB3L2 regulome within TETRAMER (Cholley et al; 2018), leading to the reconstruction of a gene regulatory network (GRN) composed by 15,148 nodes (genes) and 128,394 transcription regulation connections (edges). This has been performed when interrogating the master transcription regulatory performance for the 64 AD-specific TFs found in this study plus the homodimer components ATF4, CREB3L2 and the aberrant heterodimer ATF4-CREB3L2. **(D)** Left panel: Ranked TFs (64 + 3) based on their capacity to activate the AD-specific differentially-expressed genes (Aguzzoli Heberle et al. 2025). Master TF regulatory index per TF is displayed in blue, while the regulatory index obtained after edges randomization is displayed in gray (shaded area: Standard deviation confidence interval. Solid gray line: mean regulatory index assessed from 10 randomization processes). Right panel: TF regulatory index confidence (P-value). Green dashed line delimitates the most confident master TFs being part of the ATF4- CREB3L2 regulome (TF regulatory index >60%; P-value <0.2). Notice that only five factors are not part of this regulome, among them KLF8. **(E)** Model recapitulating the capacity of Amyloid-beta aggregates to promote the emergence of the aberrant CREB3L2-ATF4 heterodimer TF and its direct transcriptional regulation capacity over the various TFs discovered as AD-specific in human BORGs in this study and confirmed as deregulated within AD patients’ samples cohort datasets. Master TFs found as up-regulated during the cortical analysis of AD BORGs are displayed in blue.

By interrogating the chimeric ChIP-sequencing data generated by Gouveia Roque et al. we have found that several of the AD-specific master TFs revealed in APP-swe BORGs in this study are indeed controlled by the aberrant CREB3L2-ATF4 heterodimer. As illustrated in **Figure 5B**, CREB3L2-ATF4 immunoprecipitated by either HA or V5 tag pulldown appeared enriched in the promoter of KLF5, ASHL1, CREBRF, BCL11B, SATB2, TRPS1, FOXP1 and even ZNF565. Indeed, in addition to being specifically up-regulated in human brain organoids harboring AD-related genetic mutations, several of these TFs were previously linked to AD. For instance, protein levels of KLF5 have been shown to be significantly increased in both AD patients’ samples and APP/PS1 mice, and this behavior has been shown to accelerate APP amyloidogenic metabolism, notably by promoting Amyloid beta synthesis via BACE1 (Wang et al. 2022). As part of the AD-specific regulome reconstitution we have found that KLF5 directly controls the expression of ESR1 (encoding for the estrogen receptor and early on associated to AD notably via the detection of polymorphisms considered as AD risk factors (Luckhaus et Sand 2007; Ma et al. 2009)); RUNX2 (its over-expression has been shown to impair neurogenesis and oligodendrogenesis in cells harboring AD-related mutations (Nakatsu et al. 2023)); or TSHZ3 ( a TF known to regulate neuronal development, which has been also correlated to the progression of AD (Kajiwara et al. 2009)) among other factors.

The zinc protein transcription factor BCL11B - known to work as both transcriptional activator or suppressor - has been recently retrieved within a genome wide association study covering 41 families with hereditary late onset AD (Lennon et al. 2017). The Forkhead Box P1 (FOXP1) transcription factor has been shown to control the expression of the AD risk factor SORL1 (Fu et al. 2025). Finally, the zinc finger transcription factor ZNF565, known to be expressed in both the male reproductive system and the nervous system, has been highlighted as part of a study focused on human hippocampal neurogenesis in the adult and notably in AD patients (Disouky et al. 2026).

With the aim of providing a more comprehensive analysis of the influence of the ATF4- CREB3L2 heterodimer in the transcription regulation of the master TFs found in our study, we have integrated the ATF4-CREB3L2 regulome with the temporal transcriptome assessed from BORGs harboring the APP-swe mutation. In addition to evaluate bulk transcriptomes issued from samples collected at day 20 till day 120, we have also introduced in this analysis a pseudo-bulk transcriptome established from the spatially-resolved transcriptomics profiling performed in APP-swe BORGs collected at day 150. This integrative analysis, performed with TETRAMER (Cholley et al. 2018), allowed to reconstruct a gene regulatory network composed by 15,148 nodes (genes) interconnected by 128,394 transcription regulation relationships (**Figure 5C**). This reconstructed GRN has been used as template for interrogating the master regulatory capacity of all 64 TFs previously associated to AD (**Figure 4**), complemented by the TFs ATF4, CREB3L2 and the ATF4-CREB3L2 heterodimer. As illustrated in **Figure 5D**, the ATF4-CREB3L2 heterodimer presented the highest regulatory index (90,7%), indicating that the regulome associated to this aberrant heterodimer transcriptionally regulate most of the differentially expressed genes assessed during the 150 days of differentiation of BORGs harboring the APP-swe mutation. Furthermore, other TFs driven by ATF4-CREB3L2, like TRPS1, CREBRF, SATB2, KLF5, ZNF565, FOXP1 or ASH1L presented highly confident regulatory indexes (**Figure 5D**). Among all interrogated TF, RUNX3, PRDM1, HDAC4, KLF8 and ATF4 presented rather low regulatory indexes and a rather weak confidence (TF regulatory index <40%; P-value>0.6); suggesting that these factors are not driven by the ATF4-CREB3L2 regulome.

Based on these findings, in addition to the different TFs retrieved as controlled by the CREB3L2-ATF4 aberrant heterodimer, we have retained as part of the integrative regulome model other master TFs as key players of the AD-specific regulome (**Figure 5E**). Notably, the Krüppel-like factor 8 (KLF8) – which appeared up-regulated in AD-related human BORGs in this study, has been previously linked to Alzheimer’s disease in a rat model via its capacity to regulate the Wnt/Beta-catenin signaling pathway(Yi et al. 2014). Considering that KLF8 has been described as a transcriptional repressor, and that it directly interacts with the promoter of KLF5, it is tempting to speculate for an over-expression of KLF8 in an AD context as a mechanism aiming to tune-down the induction of KLF5 driven by the amyloid-beta driven CREB3L2-ATF4 aberrant heterodimer. This being said, other studies described KLF8 as a transcription activator, thus opening the way to a direct transcription activation mechanism of the various AD-specific TFs retrieved within the reconstituted regulome (Krüppel-like factor 8 activates the transcription of C-X-C cytokine receptor type 4 to promote breast cancer cell invasion, transendothelial migration and metastasis s. d.; Urvalek et al. 2008).

Another factor that is not directly controlled by the CREB3L2-ATF4 heterodimer is the hypoxia-inducible transcription factor (HIF3A), which has been highlighted as a key player within a study focused on evaluating the oxidative stress as a hallmark of AD. Indeed, HIF3A mediates an aberrant oxidative stress response in AD via the transcriptional induction of the factor TXNIP (Xue et al. 2026). Similarly, the zinc finger transcription factor TSHZ3 has been also correlated to AD, notably by the discovery of a rare variant in this gene leading to Amyloid beta aberrant processing (Kajiwara et al. 2009; Louwersheimer et al. 2017).

In summary, at this stage we propose a model that integrates the capacity of Beta amyloid to induce the aberrant CREB3L2-ATF4 heterodimer formation with the downstream activation of the AD-specific regulome that we have reconstituted in this study (**Figure 5E**).

## DISCUSSION

In this study we have reconstituted the regulome that is setup in presence of AD-related genetic mutations and confirmed their prevalence in human patients’ samples. Among the 64 top-confident master transcription factors, both KLF8 and KLF5 appeared as major players, notably due to the presence of such binding motifs in roughly half of the AD-specific master TFs. While the bulk transcriptomics data assessed on BORGs tend to suggest that KLF5 is not differentially expressed in the APP-swe genetic background (**Figure 3H** **& Supplementary Figure S8**), the spatially-resolved transcriptomics (SrT) data demonstrated that KLF5 is significantly up-regulated in APP-swe BORGs relative to the corresponding isogenic wild-type counterpart (**Supplementary Figure S11E**). This discrepancy could be explained by the differences on chronological age between the analyzed samples (5 months for SrT; up to 4 months for bulk transcriptomics) and notably influenced by the delayed progression in maturation (**Figure 1** & Figure 2).

KLF5 has been found as a direct player influencing AD progression in mice models (APPswe/PS1De9), notably via its positive regulation of BACE1, further suggesting that the upregulation of KLF5 is a critical element in AD(Wang et al. 2022). Similarly, KLF8 has been previously liked to the progression of AD notably based on its capacity to regulate the Beta-catenin / Wnt signalling pathways (Yi et al. 2014)). As consequence, the deregulation of these two factors is expected to have major impact based on the AD-specific regulome revealed in this study. This being said, the question that remained open is the link between the deregulated gene expression changes observed in AD and the pathophysiology of the disease. Importantly, Gouveia Roque and colleagues demonstrated recently that the exposure of neurons to oligomerized Amyloid Beta 42 induce the aberrant heterodimerization of the CREB3L2-ATF4 transcription factors (Gouveia Roque et al. 2023). The analysis of the chimeric ChIP-sequencing data generated by Gouveia Roque et al. allowed us to confirm that several of the master TF players composing the AD-specific regulome described in this study presented CREB3L2-ATF4 enriched at their promoters; among them KLF5 (**Figure 5**). This finding clearly supports a model in which the beta-amyloid plaques could directly influence the setup of an aberrant regulome, passing via the installation of the CREB3L2-ATF4 heterodimer leading to the activation of the regulome highlighted in this study.

## MATERIALS AND METHODS

### Cell culture of iPSCs

Human induced pluripotent stem cells (iPSCs) were obtained from the New York Stem Cell Foundation (NYSCF). Notably the wild-type parental line 7889SA and the knock-in constructs APP Swe/Swe (CO0002-01-CS-004 / BN0013), PSEN1 M146V/M146V (CO0002-01-CS-001) and Double Knock-In Swe/Swe + M146V/M146V (BN0002) were used in this study. This constructs were initially described by (Kwart et al. 2019). IPSCs were passaged and maintained using standard feeder-free culture protocols. Briefly, iPSCs were grown on Matrigel-coated plates (Corning) in mTeSR1 (STEMCELL Technologies) and passaged each two days. Cells at 70-80% confluency are split in presence of ReLeSR (STEMCELL Technologies).

### Brain organoid cultures

Brain organoid cultures are based on the original protocol described by Roseborck et al (Rosebrock et al. 2022) relying on the use of a triple-inhibitors action (Dual SMAD, TGFB and WNT inhibition). Briefly, 9,000 hIPSCs are deposited on a U-bottom 96 well plate in presence of the embryoid body (EB) formation medium supplemented by Rock Inhibitor (50uM), bFGF (4ng/ml), SB431542 (10µM), LDN193189 (100nM) et XAV939 (3,3µM). EBs are refreshed each two days by replacing half of the medium. After 7 days EBs medium is replaced by Neural induction medium (NIM) containing the neuro-induction molecules SB431542 (10µM), LDN193189 (100nM) et XAV939 (3,3µM). NIM medium is refreshed at day 10 and at day 12 the EBs are embedded in Matrigel growth factor reduced droplets. Solidified droplets at 37°C are transferred into 6-well plates containing expansion medium (DMEM/F12, NB, Glutamax supplement, MEM-NEAA, N2, B27 without Vitamin A, Insulin, Beta-mercaptoEthanol). Expansion medium is kept until day 16, then replaced by maturation medium presenting B27+VitaminA. BORGs are cultured under agitation in 6-well plates over several months in presence of maturation medium (refreshed each 4 days). Three different brain organoid kinetics were produced for WT control and APP Swedish samples; and two different kinetics for PSEN1-M146V and for the APP-Swedish / PSEN1-M146V double mutant.

### Beta-amyloid quantification

For ELISA, three BORGs cultured during 120 days were dissociated together in presence of SDS lysis buffer. Total protein levels were quantified with Bradford reagent. 20ug of total protein has been used for ELISA quantification targeting both the Amyloid Beta42 (thermo Fisher Scientific khb3544) and the Amyloid Beta40 peptides (thermo Fisher Scientific KHB3481). ELISA quantification has been performed at least in technical duplicates. These ELISA measurements were performed in samples issued from three different brain organoid kinetics for WT control and APP Swedish samples; and from two different kinetics for the APP-Swedish / PSEN1-M146V double mutant.

### Immunohistochemical staining

BORG tissue sections were permeabilized (Triton 0.1% in PBS; 15 minutes at room temperature) and blocked (0.1% Triton and 1% BSA in PBS during 1 h at room temperature). Sections were washed 3 × 5 minutes in permeabilization buffer, then incubated with the primary antibodies: anti-β III tubulin/anti-TUBB3 (ab14545; Abcam); anti-MAP2 (ab32454; Abcam); anti-GFAP (ab10062; Abcam); anti-Beta-42 (Cell Signaling; 8243); anti-SOX2 (ab97959; Abcam). After 1 h of incubation at room temperature (or overnight at +4°C), sections were washed 3 × 10 min with permeabilization buffer followed by incubation with a secondary antibody (donkey anti-mouse IgG [H + L] antibody Alexa 555; Invitrogen A- 31570; donkey anti-rabbit IgG [H + L] antibody Alexa 488; Invitrogen A-21206) and/or DAPI (D3571; Invitrogen). After 1 h at room temperature, sections were washed for 3 × 10 min in permeabilization buffer twice with Milli-Q water and finally mounted on microscope slides.

### RNA isolation, cDNA synthesis and library preparation for bulk transcriptomics

Total RNA has extracted from at least three BORGs using RNeasy Mini kit (Qiagen 74104). RNA-sequencing libraries were performed with the NEBNext Ultra II RNA Library Prep Kit for Illumina (E7770). Libraries were sequenced within the French National Sequencing Center, Genoscope (150-nt pair-end sequencing; NovaSeq Illumina; >25 million reads per library).

### Primary Bulk transcriptomics analysis

Fastq files were mapped to the human reference genome hg19 using Bowtie 2.1.0 under default parameters. Mapped reads were associated with known genes with featureCounts. RNA-seq analyses were done with the DESeq2 R package. Heatmap matrix display was generated with MeV 4.9.0. Gene Ontology analyses were performed with EnrichR: https://maayanlab.cloud/Enrichr/. Neural cell lineage enrichment analyses were performed by computing a hypergeometric probability test from the differential expression data and a target gene set using customized “R” scripts. Differentially expressed genes were challenged into a gene-set collection concerning the different neural cell lineage markers, obtained by manual curation from different publications (**Supplementary Table S1**).

### Brain organoids cryo-sectioning and Spatially-resolved transcriptomics (SrT)

BORGs were fixed with 4% paraformaldehyde (over-night) and incubated in Sucrose 30% (over-night). Several BORGs were placed in plastic molds, covered with Optimal Cutting Temperature (OCT) embedding medium and frozen prior being cryo-sectioned (Cryo-star).

Tissue sections of 10 microns of thickness were placed on top of in-house manufactured double-barcoded DNA arrays. Briefly, these arrays are composed by 2,048 DNA probes presenting a poly-T sequence at the 3’-end for capturing messenger RNA and two molecular barcodes corresponding to their physical coordinates (X, Y) within the array (Lozachmeur et al. 2023).

During tissue cryo-sectioning, DNA arrays were kept inside the cryostat chamber to preserve RNA integrity. DNA array slides are briefly warmed by placing a finger in the back region where tissue should be deposited to allow tissue attachment. Deposited tissue sections were processed for library preparation as described in our in-house protocol (Lozachmeur et al. 2023). SrT libraries were sequenced within the French National Sequencing Center, Genoscope (150-nt pair-end sequencing; NovaSeq Illumina; >200 million reads per library).

### Spatially-resolved transcriptomics processing

Primary analysis has been performed with our in-house developed tool SysISTD (SysFate Illumina Spatial transcrip- tomics Demultiplexer: https://github.com/SysFate/SysISTD). As outcome, SysISTD generates a matrix presenting read counts per gene ID in rows and spatial coordinates in columns. To focus the downstream analysis to the physical positions corresponding to the analyzed tissue, we used an in- house R script taking as entry an image of the DNA array scanned with the TRICT filter, revealing the presence of the fiducial borders introduced during the manufacturing of the DNA arrays.

Focused matrices were processed with our in-house Spatial omics depixelation tool allowing to pass from matrices of 64x64 pixels till 128x128 pixels (Mendoza-Parra, Duvina, et Galindo-Albarrán 2025). Depixelated matrices were quantile normalized and used for computing the differential gene expression levels per pixel (relative to the average levels across the whole tissue) within our previously described tool MULTILAYER (Moehlin, Koshy, et al. 2021; Moehlin, Mollet, et al. 2021).

### Temporal gene co-expression maps stratification and GRN reconstruction

Temporal gene expression data were stratified in multiple gene co-expression paths with DREM 2.0 (Schulz et al. 2012). Gene regulatory networks were reconstructed with TETRAMER (Cholley et al. 2018). As entry-point TETRAMER incorporated the differential gene expression data generated per BORG and a collection of TF-TG relationships issued from the CellNet collection (available within TETRAMER). CENTRIMO motif analysis has been performed within MEME suite 5.5.9 https://meme-suite.org/meme/tools/meme.

Top-ranked TFs were used for building the AD-specific co-regulatory network within Cytoscape 3.4.0.

### Comparison with human AD patient cohorts

Alzheimer’s disease (AD)-associated transcriptional signatures were obtained from precomputed bulk RNA-sequencing differential-expression results available through Alzheimer DataLENS (Noori et al. 2024), an open-data platform that consistently processes public AD omics datasets, including datasets from the AMP-AD Knowledge Portal. We selected 18 contrasts from three independent postmortem brain cohorts: the Mayo Clinic cohort (Allen et al. 2016) comprising cerebellum (CBE) and temporal cortex samples (TCX); the Mount Sinai cohort comprising frontal pole (FP), inferior frontal gyrus (IFG), parahippocampal gyrus (PHG), and superior temporal gyrus (STG) samples; and ROSMAP (Mostafavi et al. 2018) comprising prefrontal cortex (PFC) samples. The selected contrasts represented AD-associated changes defined by clinical diagnosis or by clinical and neuropathological measures, including Braak neurofibrillary tangle stage, CERAD neuritic plaque score, Clinical Dementia Rating (CDR), and Mini-Mental State Examination (MMSE) score.

For each comparison, AD-associated signature was composed of genes which were considered AD-upregulated if they had a log2 fold change greater than 0.25 and a Benjamini– Hochberg-adjusted P value less than 0.05. Duplicate gene symbols were removed, retaining the highest-ranked occurrence. For signatures containing more than 1,000 differentially expressed genes, the top 1,000 genes showing the largest changes were retained; signatures containing fewer than 20 genes were excluded from subsequent analyses.

To assess the spatial enrichment of these AD-associated signatures in wild-type and APP-Swe BORGs, gene set enrichment analysis was performed independently for each pixel using the fgsea (v1.30.0) package. Within each pixel, genes were ranked in descending order according to their pixel-specific differential values, while the AD-associated gene signatures were used as the gene sets. Pixels-specific differential values correspond to gene normalized counts per pixel relative to the gene average counts across all pixels, as performed by MULTILAYER(Moehlin, Koshy, et al. 2021; Moehlin, Mollet, et al. 2021). The multilevel FGSEA algorithm was used by default; when it failed or returned missing normalized enrichment scores, the permutation-based implementation was applied with 1,000 permutations. The enrichment results were combined across all successfully processed pixels. **Reanalysis of ChIP-sequencing dataset assessed from CREB3L2-ATF4 heterodimer** Chimeric ChIP-sequencing data generated by Gouveia Roque et al. were interrogated for the presence of CREB3L2-ATF4 heterodimer binding sites at the promoter regions of the AD-specific master TFs revealed in APP-swe BORGs by using the UCSC tracks made publicly available by the authors: https://genome.ucsc.edu/s/CGRoque/GouveiaRoque_ChIPmera

CREB3L2-ATF4 ChIP-seq profiles were downloaded from GEO repository (GSE147205) in bigwig format, then converted into bedgraph, followed by peak calling processing with MACS2. Obtained peaks were annotated to their proximal gene (hg38 assembly) and used for defining the CREB3L2-ATF4 regulome by conserving genes in a proximal distance of 5000 bases. CREB3L2-ATF4 regulome has been used in TETRAMER together with the differentially expressed genes obtained in APP-swe BORGs issued from day 20, 30, 60, 90 and 120. In addition, a pseudo-bulk transcriptome profiling has been obtained from APP-swe BORGs analyzed by spatial transcriptomics (day 150), by selecting the most up-regulated levels per gene across all analyzed pixels. TETRAMER processing has been performed by interrogating as starting genes the set of 64 AD-specific TFs found from the comparison between BORGs and AD patients’ data and complemented with ATF4, CREB3L2 and the CREB3L2-ATF4 heterodimer. As final nodes, the differentially expressed genes associated to AD patients’ data provided by (Aguzzoli Heberle et al. 2025).

## ACKNOWLEDGMENTS

We thank all members of the team SysFate for contributing to the discussion of this project and the Genoscope sequencing platform for their technical support.

## AUTHOR CONTRIBUTIONS

Antoine Aubert (Investigation, Data curation, Software, Formal analysis), Anne-Claire Comby (Formal analysis, Investigation, Methodology), Aude Bramoulle (Formal analysis, Investigation, Methodology), Maria Grazia Mendoza-Ferri (Resources, Methodology), Peggy Azzolin (Resources, Methodology), Alice Moussy (Resources, Methodology), Huan Li (Resources, Methodology), Sudeshna Das (Supervision, Funding acquisition, revision original draft), Bruno Maria Colombo (Supervision, Funding acquisition, revision original draft) and Marco Antonio Mendoza-Parra (Conceptualization, Formal analysis, Supervision, Funding acquisition, Writing original draft).

## DECLARATION OF INTERESTS

None declared.

## FUNDING

This work was supported by the institutional bodies CEA, CNRS, Universitéd’Evry-Val d’Essonne. Furthermore, this project has been supported by the Genopole Thematic Incentive Actions funding (ATIGE-2017), the “Fondation pour la Recherche Medicale” (FRM; funding ALZ-201912009904), and the “Alzheimer’s Research foundation” (FRA; funding 2023-A-01; Bruno-Maria Colombo). Aude Bramoulle has been funded by the Institut National du Cancer (INCa: funding 2020-181). Maria Grazia Mendoza-Ferri has been funded by the Institut National du Cancer (INCa: Funding N ◦2020-124; N ◦2022- 078).

## DATA AVAILABILITY

All raw datasets generated on this study have been submitted to the NCBI Gene Expression Omnibus (GEO; http://www.ncbi.nlm.nih.gov/geo/) under accession number XXXXX

## Notes

### Competing Interest Statement

The authors have declared no competing interest.

### Summary of Updates

Figure 1 and Figure 5 were improved.

