## Supplementary_Figures for "Krüppel-like factors KLF5 & KLF8 emerge as potential master transcriptional regulators of Alzheimer’s disease, as revealed on cell fate regulomes in human brain organoids"

### Supplementary Material

**Supplementary Table S1.** Gene set collection concerning the different neural cell lineage markers.
**Supplementary Table S2.** Upregulated genes spatially overlapping with the various cortical layer regions defined by different gene markers.
**Supplementary Figure S1.** Human brain organoids size progression.
**Supplementary Figure S2.** Immunofluorescence micrograph of human brain organoids issued from human induced pluripotent stem cells harboring familial Alzheimer's disease genetic mutations and their isogenic control « WT » line targeting the markers TUBB3 and SOX2.
**Supplementary Figure S3.** Immunofluorescence micrograph of human brain organoids issued from human induced pluripotent stem cells harboring familial Alzheimer's disease genetic mutations and their isogenic control « WT » line targeting the markers TUBB3 and GFAP.
**Supplementary Figure S4.** Differential gene co-expression changes during BORGs progression.
**Supplementary Figure S5.** Differential gene expression changes between BORGs of 120 days of age.
**Supplementary Figure S6.** Temporal differential gene expression signatures during brain organoids differentiation associated to major Cell-type markers.
**Supplementary Figure S7.** Gene regulatory networks (GRNs) reconstructed for AD and WT BORGs.
**Supplementary Figure S8.** Expression of the transcription Krüppel-like factor family during BORGs differentiation.
**Supplementary Figure S9.** Pixel-level gene set enrichment analysis (GSEA) of AD-upregulated signatures retrieved from Alzheimer DataLENS in AD-related Brain organoids.
**Supplementary Figure S10.** Pixel-level gene set enrichment analysis (GSEA) of AD-upregulated signatures retrieved from Alzheimer DataLENS in WT Brain organoids.
**Supplementary Figure S11.** Spatially-resolved transcriptomics (SrT) assay performed on WT and APP-Swedish BORGs of 5 months of age.

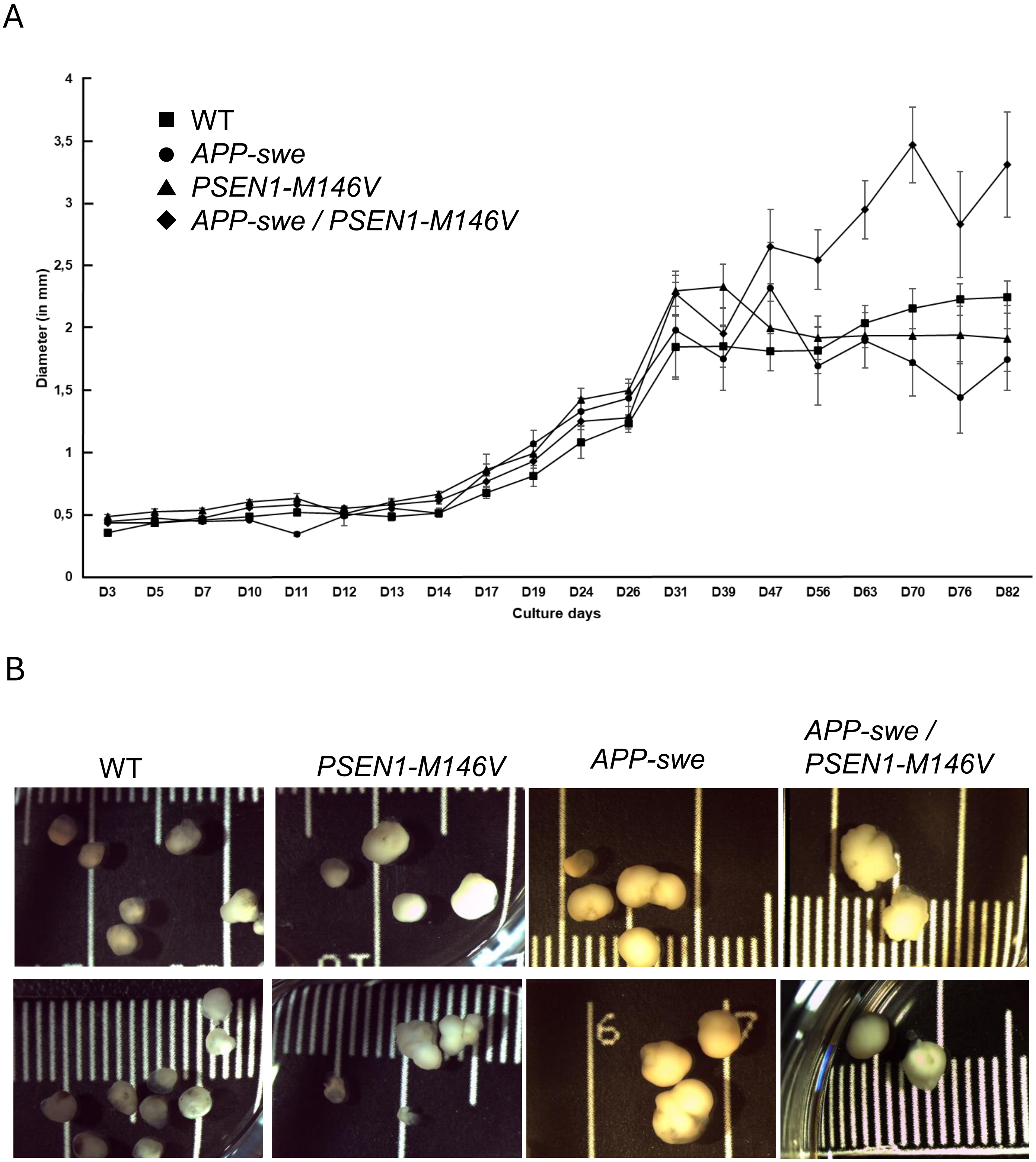


**Supplementary Figure S1. Human brain organoids size progression. (A)** Human induced pluripotent stem cells harboring the familial Alzheimer's disease genetic mutations *APP-swe*, *PSEN1-M146V,* a double mutant, as well as their isogenic parental « Wild-type » (WT) were cultured over several months under cortical brain organoid (BORG) culture conditions. During BORGs differentiation, their gain in size was monitored by bright field imaging. Notice that for all hIPS lines, BORGs size followed a similar progression during the first 40 days, reaching an average diameter size of 2 mm. Importantly, in the next 40 days BORGs size remained constant in average, with the exception of the hIPS line harboring both mutations, which managed to reach 3 mm in average. **(B)** Representative bright field images of BORGs assessed at 120 days.

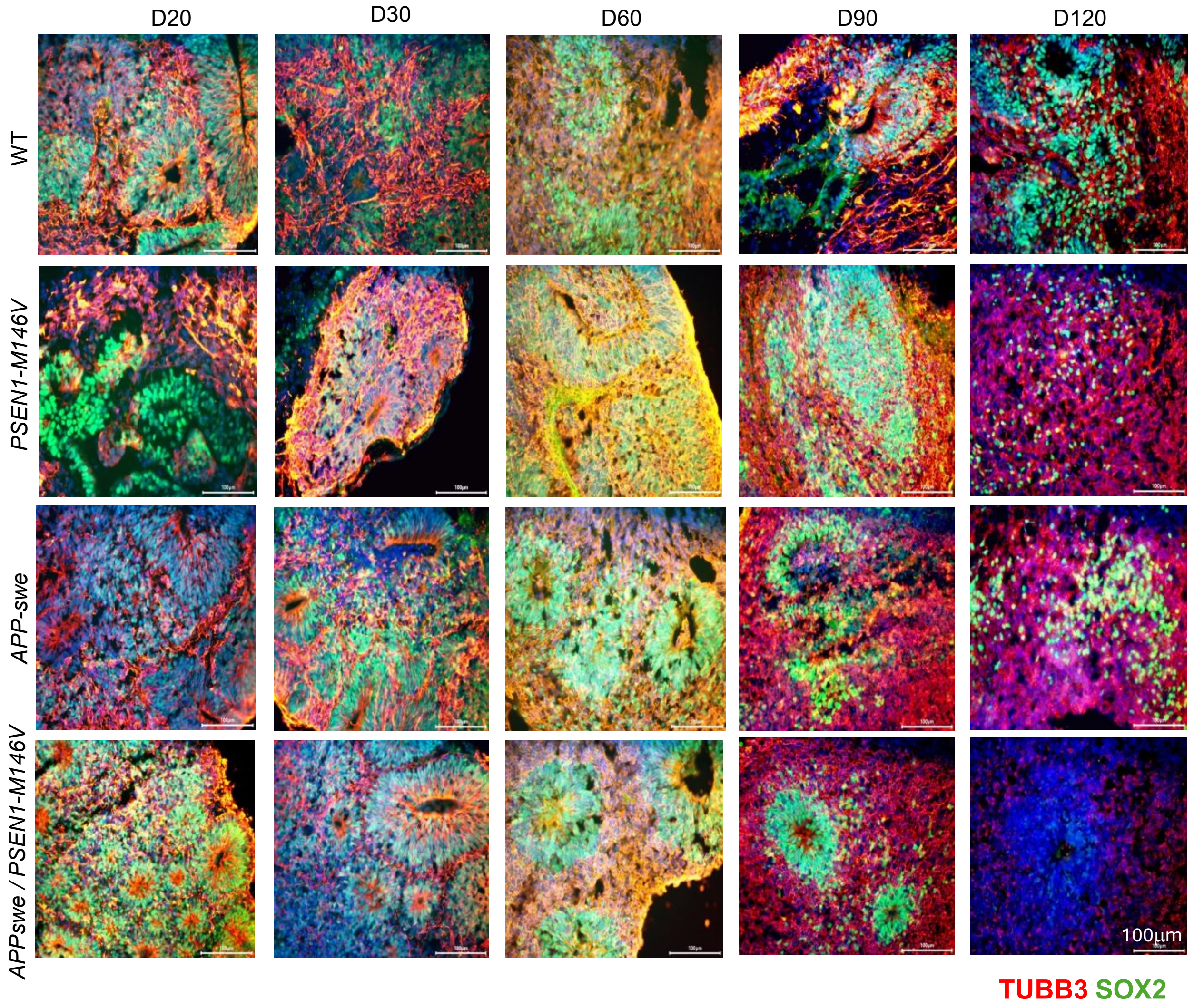


**Supplementary Figure S2. Immunofluorescence micrograph of human brain organoids issued from human induced pluripotent stem cells harboring familial Alzheimer's disease genetic mutations and their isogenic control « WT » line targeting the markers TUBB3 and SOX2.** Brain organoids collected at various time-points during their differentiation were stained for the neuronal marker TUBB3 (red) and the pluripotent neuronal stem cell marker SOX2 (green). Notice that SOX2 immunostaining reveals the presence of neural rosette structures, witnessing the presence of neural stem cell progenitors. Bar-size displayed on micrographs correspond to 100 µm.

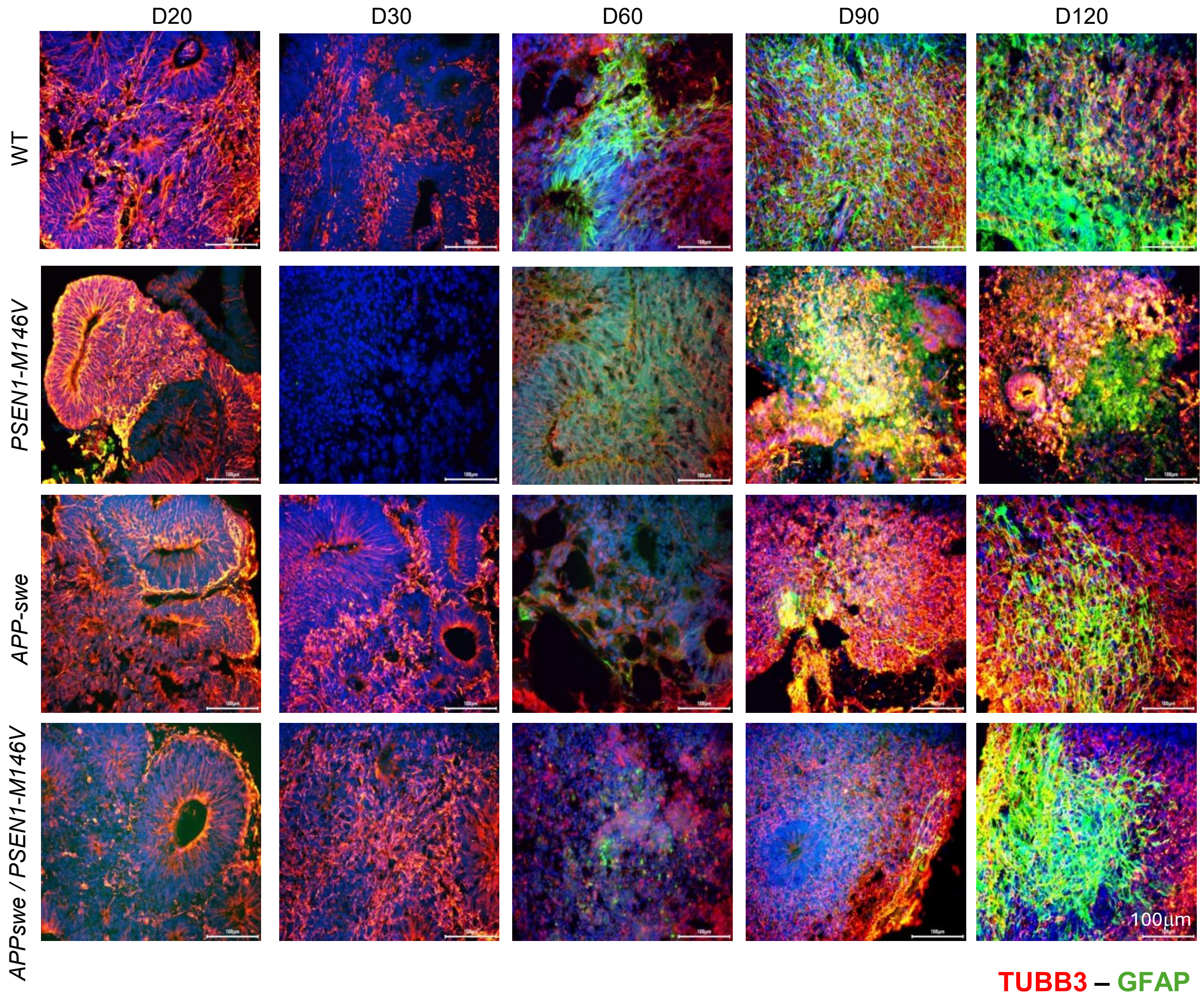


**Supplementary Figure S3. Immunofluorescence micrograph of human brain organoids issued from human induced pluripotent stem cells harboring familial Alzheimer's disease genetic mutations and their isogenic control « WT » line targeting the markers TUBB3 and GFAP.** Brain organoids collected at various time-points during their differentiation were stained for the neuronal marker TUBB3 (red) and the astrocyte marker GFAP (green). Notice that GAP-positive cells are observed from 60 days of differentiation on WT BORGs and from 90 days for those harboring the Alzheimer's disease-related mutations. Bar-size displayed on micrographs correspond to 100 μm.

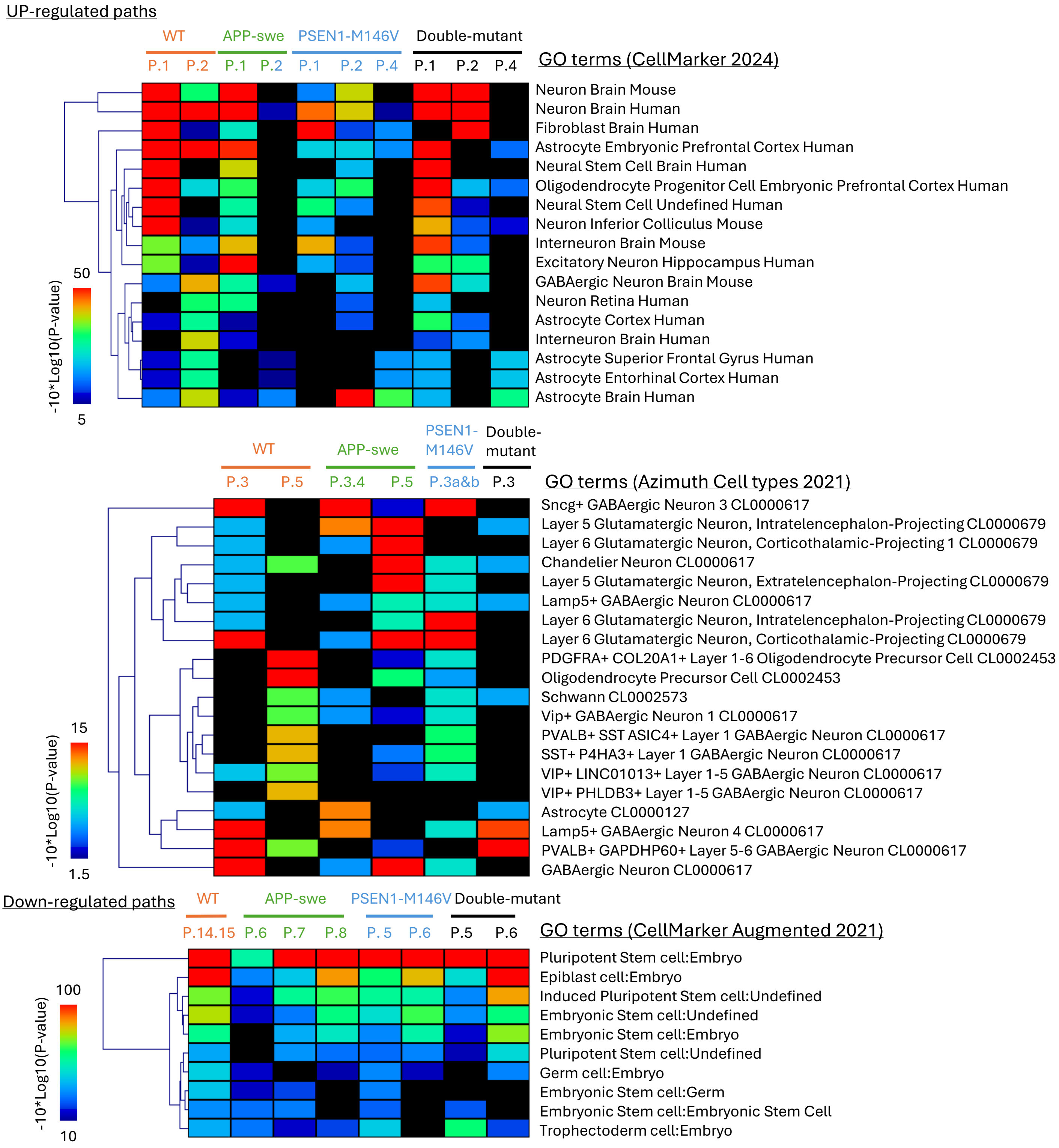


**Supplementary Figure S4. Differential gene co-expression changes during BORGs progression. (A)** Gene co-expression paths profiling assessed for BORGs harboring AD-related mutations. Co-expression paths presenting differential gene expression levels higher than 2-fold (Log2) are displayed. **(B)** GO terms enrichment confidence assessed on WT BORGs and its comparison with those assessed on AD-related BORGs.

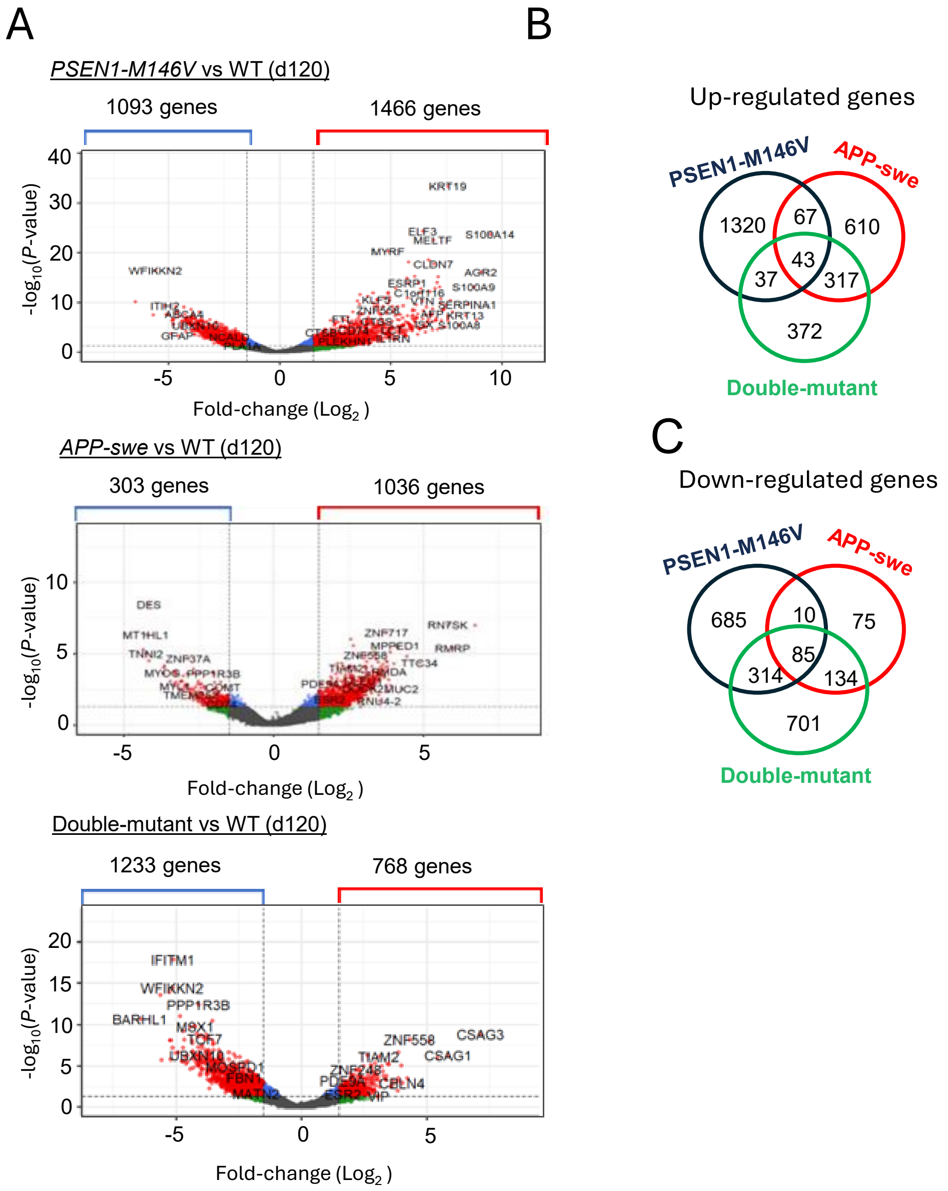


**Supplementary Figure S5. Differential gene expression changes between BORGs of 120 days of age. (A)** Volcano plot displays revealing the differential gene expression changes between BORGs harboring AD-related mutations and their isogenic WT counterpart after 120 days of differentiation. **(B)** Venn-diagram displaying the common and specific up-regulated genes between BORGs harboring the AD-related mutations relative to the WT control line. Notice the low number of common genes arguing for major discrepancies on BORGs progression. **(C)** Same as (B) but performed for down-regulated genes.

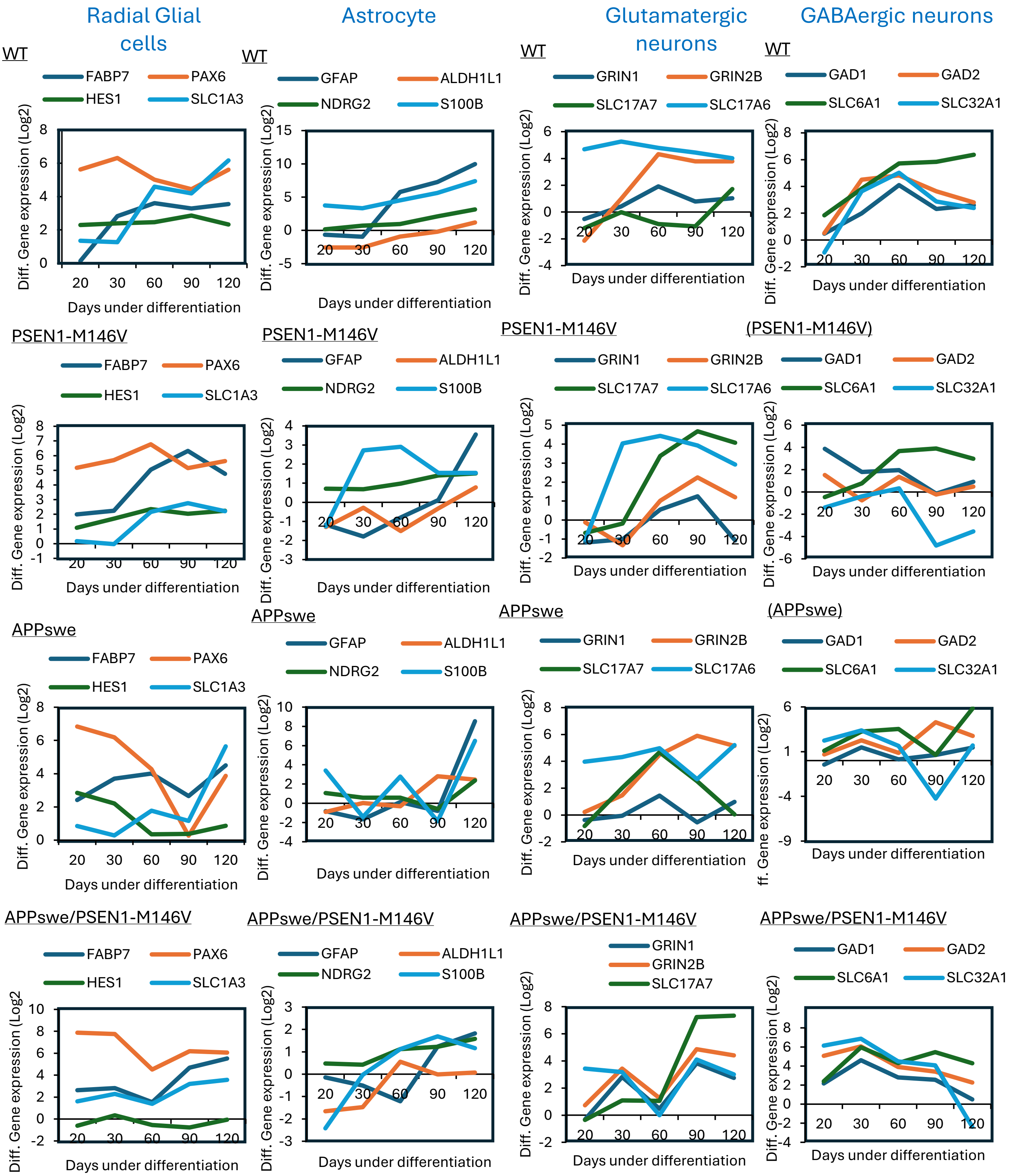


**Supplementary Figure S6. Temporal differential gene expression signatures during brain organoids differentiation associated to major Celltype markers.**

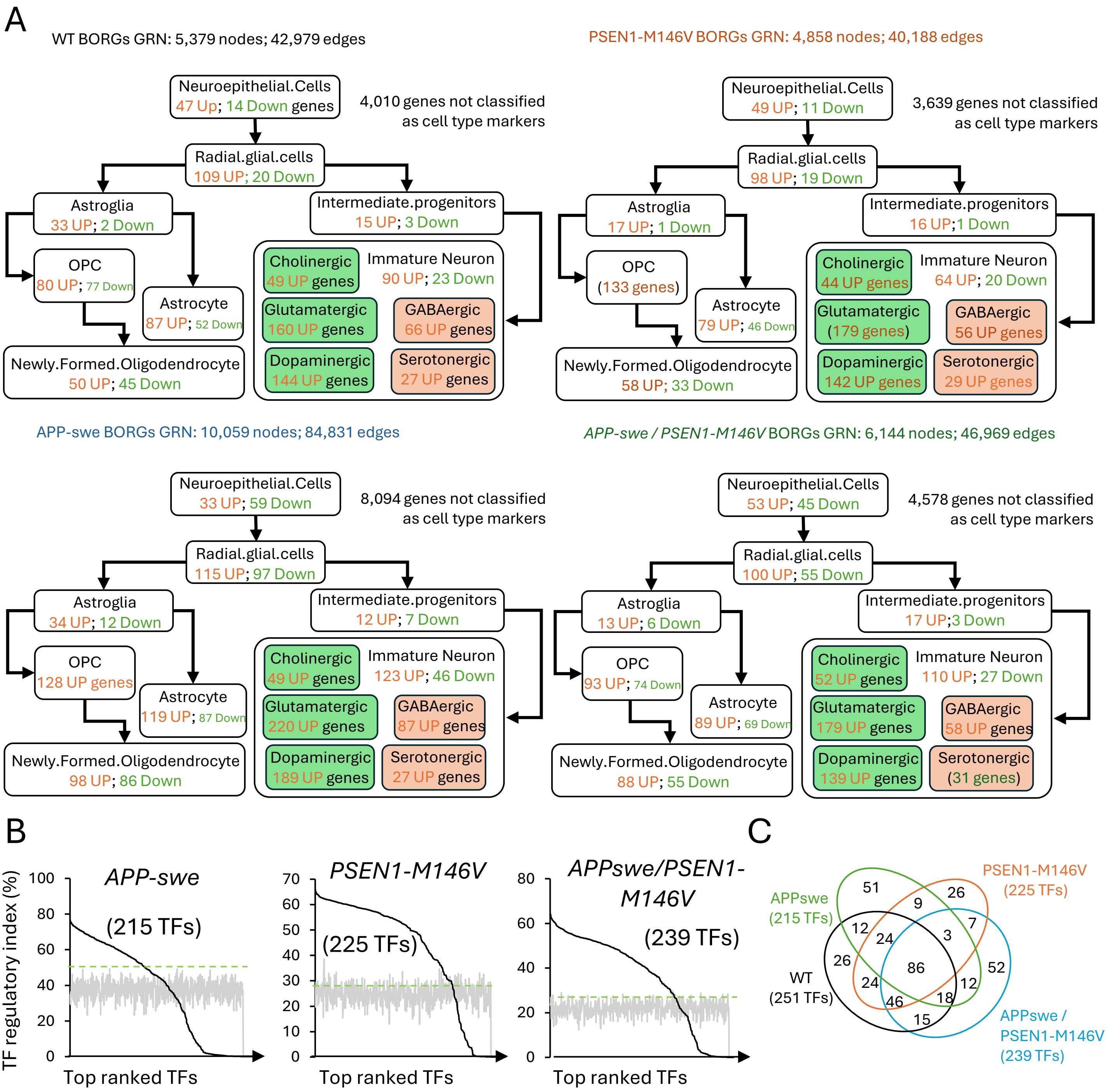


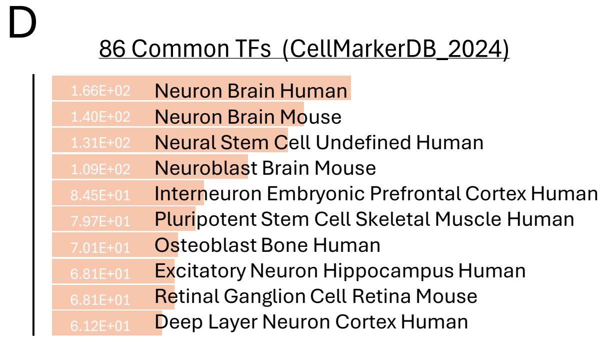


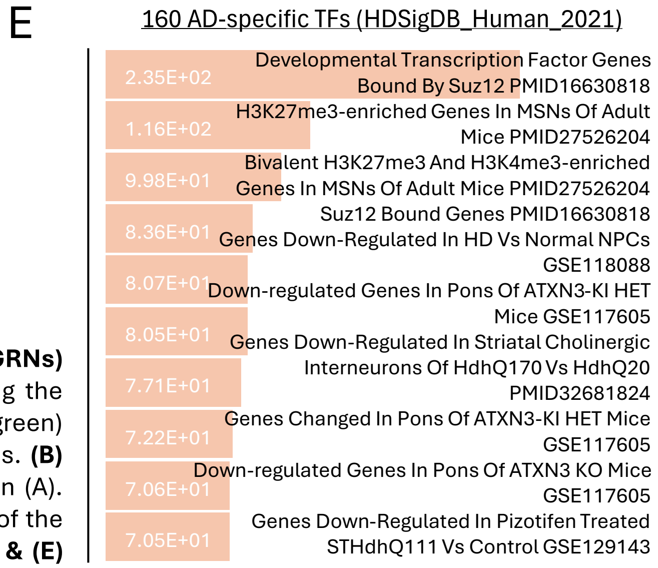


**Supplementary Figure S7. Gene regulatory networks (GRNs) reconstructed for AD and WT BORGs. (A)** Scheme displaying the presence of Up-regulated (orange) or down-regulated genes (green) associated to each of the indicated cell-types within the GRNs. **(B)** Master TFs identified by TETRAMER over the GRNs displayed in (A). **(C)** Venn-diagramme comparing the master TFs found in each of the AD-related BORG samples with their isogenic WT control. **(D) & (E)** Gene Ontology enrichment anlaysis performed on the 86 common as well as th e160 AD-specific TFs displayed in (C).

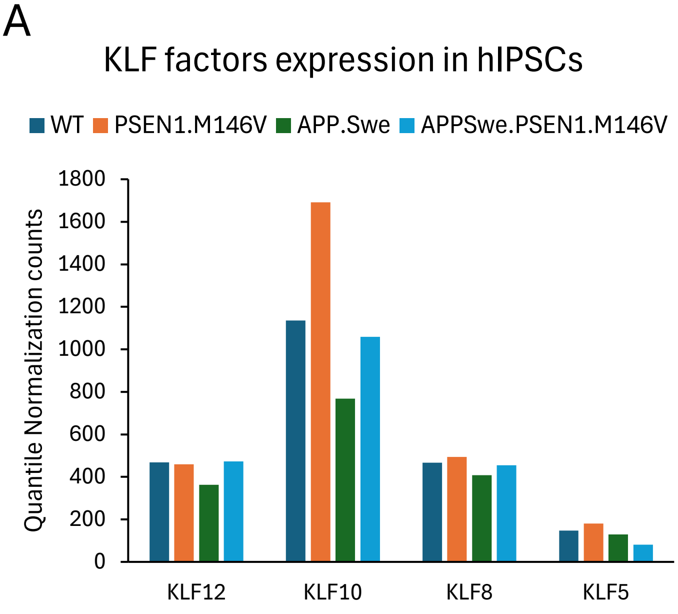


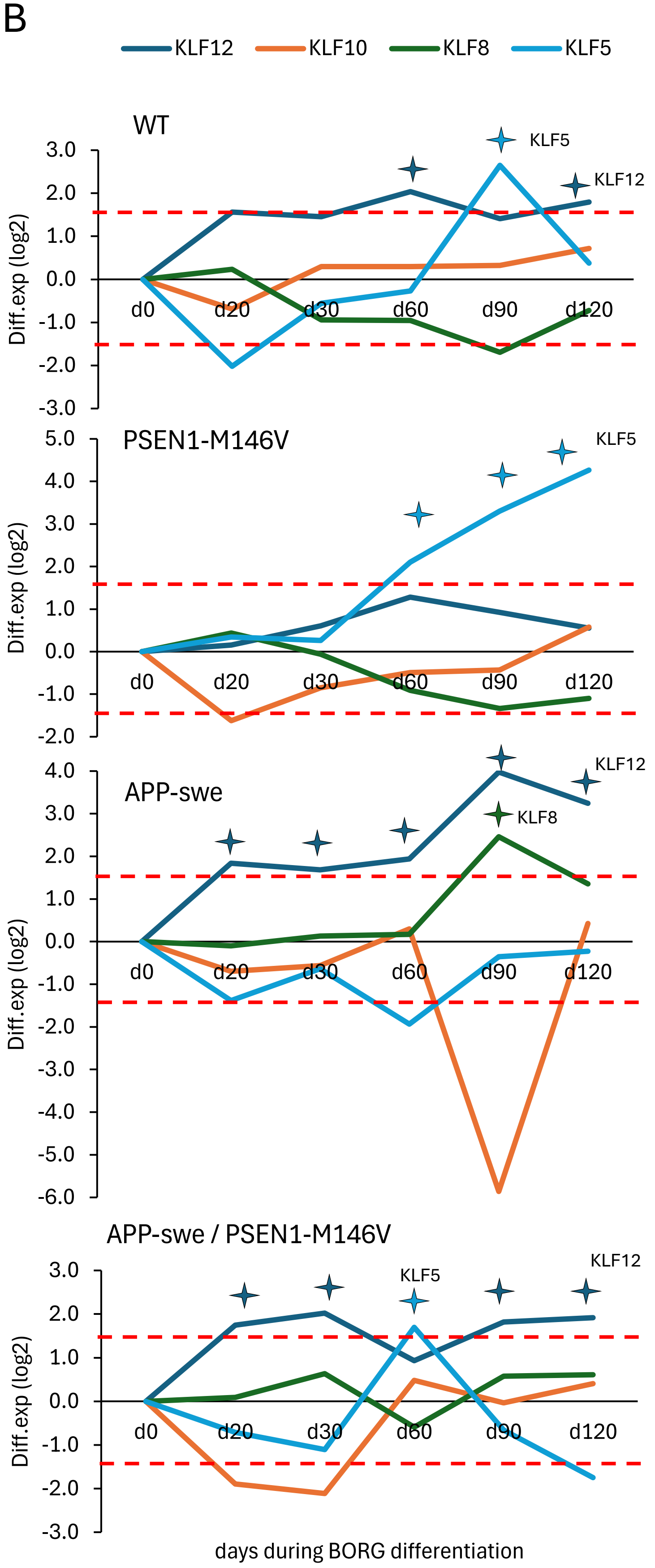


**Supplementary Figure S8. Expression of the transcription Krüppel-like factor family during BORGs differentiation. (A)** KLF transcription levels retrieved in WT human induced pluripotent stem cells or in those harbouring the AD-related mutations. **(B)** Differential expression of the indicated KLF transcription factors during BORGs progression. Over-expressed TFs (LFC>1.5) is labelled with a star.

MayoBB_TCX_CPM_ClinicalDiagnosis_AD-NCI

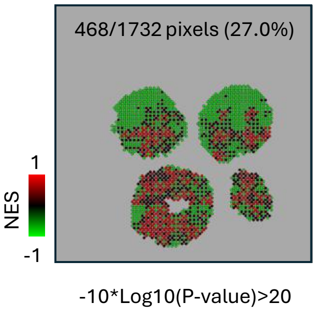


276 subjects : Alzheimer's disease (AD), N=82, progressive supranuclear palsy (PSP), N=84, pathologic aging (PA), N=30, and control (CON), N=80. Allen, Mariet et al; Scientific data vol. 3 160089. 11 Oct. 2016. **Brain region: Temporal Cortex (TCX).** Differential Gene expression analysis from Normalized RNA-seq data: AD vs No cognitive impairment (NCI) control samples.

MayoBB_CBE_CPM_Clinical Diagnosis_AD-NCI

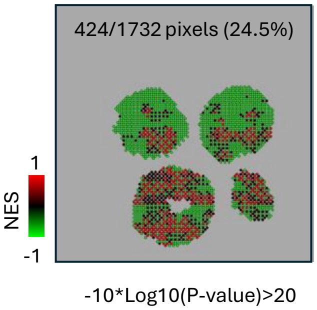


278 subjects: Alzheimer's disease (AD), N=86, progressive supranuclear palsy (PSP), N=84, pathologic aging (PA), N=28, and control (CON), N=80. Allen, Mariet et al; Scientific data vol. 3 160089. 11 Oct. 2016. **Brain region: Cerebellum (CBE).** Differential Gene expression analysis from Normalized RNA-seq data: AD vs No cognitive impairment (NCI) control samples.

MSBB_PHG_CPM_CDR_AD-NCI

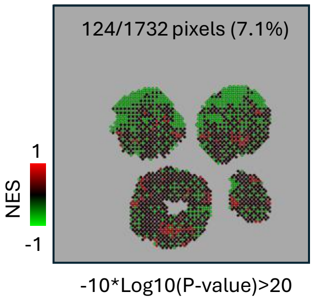


244 subjects: Alzheimer's disease (AD), N=176, No cognitive impairment (NCI), N=68. **Clinical Dementia Rating (CDR): AD vs NCI**. Wang, Minghui et al. "The Mount Sinai cohort of large-scale genomic, transcriptomic and proteomic data in Alzheimer's disease." Scientific data vol. 5 180185. 11 Sep. 2018. **Brain region : Parahippocampal Gyrus (PHG).** Differential Gene expression analysis from Normalized RNA-seq data: AD vs No cognitive impairment (NCI) control samples.

MSBB_PHG_CPM_Braak_B3-B1

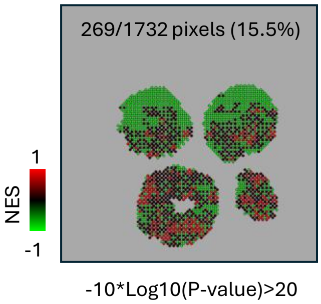


244 subjects: Alzheimer's disease (AD), N=176, No cognitive impairment (NCI), N=68. **Braak Stage** (neuropathological score focused on tau neurofibrillary tangles) B3 (stage V/VI; neocortical) vs B1 (stage 0; normal, I/II (hippocampal) . Wang, Minghui et al. Scientific data vol. 5 180185. 11 Sep. 2018. **Brain region : Parahippocampal Gyrus (PHG).** Differential Gene expression analysis from Normalized RNA-seq data between samples labeled as Braak stage B3 vs B1.

MSBB_PHG_CPM_CERAD_C3-C0

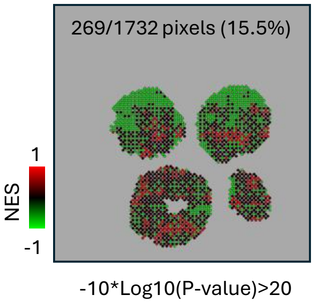


244 subjects: Alzheimer's disease (AD), N=176, No cognitive impairment (NCI), N=68. **CERAD stage** (Neuropathological score focused on the frequency of Beta-amyloid neuritic plaques) C3 (Frequent/Definite AD) vs C0 (None / Not AD). Wang, Minghui et al. "The Mount Sinai cohort of large-scale genomic, transcriptomic and proteomic data in Alzheimer's disease." Scientific data vol. 5 180185. 11 Sep. 2018. **Brain region : Parahippocampal Gyrus (PHG).** Differential Gene expression analysis from Normalized RNA-seq data between samples labelled as CERAD C3 vs C0.

ROSMAP_PFC_FPKM_Braak_B3-B1

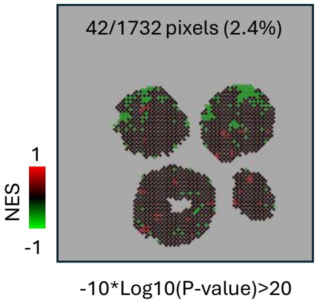


1102 subjects: Alzheimer's disease (AD), N = 396, No Cognitive Impairment (NCI), N = 351, Mild Cognitive Impairment(MCI), N = 255, AD + other, N = 56, Non- AD Dementia, N = 23, MCI + other, N = 16. **Braak Stage** (score focused on tau neurofibrillary tangles) B3 (stage V/VI; neocortical) vs B1 (stage 0; normal, I/II (hippocampal). Mostafavi, Sara et al. "A molecular network of the aging human brain provides insights into the pathology and cognitive decline of Alzheimer's disease." Nature neuroscience vol. 21,6 (2018): 811-819. **Brain region : Prefrontal Cortex (PFC).** Differential Gene expression analysis from pre-normalized expression data (FPKM calls) between samples labeled as Braak stage B3 vs B1.

**Supplementary Figure S9. Pixel-level gene set enrichment analysis (GSEA) of AD-upregulated signatures retrieved from Alzheimer DataLENS in AD-related Brain organoids.** Spatially-resolved transcriptomics (SrT) data assessed from 5-months old APP-Swedish brain organoids has been interrogated for enrichment of AD-upregulated signatures derived from human AD patient cohorts. These signatures were defined using different disease-stratification strategies, including Clinical Dementia Rating (CDR), Braak stage, and CERAD score, across multiple brain regions and data collections. Each AD-upregulated signature was used as a gene set for pixel-level GSEA of the SrT data. GSEA outcomes were first P-value filtered (-10*Log10(P-value)>20), then expressed as normalized enrichment score (NES) per pixel region (Heatmap). For each of the displayed maps the source of the data samples used as gene set, the analyzed brain region, as well as the AD scoring strategy is described, as retrieved in the DataLENS database. The analyzed APP-swe 5 months BORGs did not present a significant enrichment (-10*Log10(P-value)>20) for genes associated to transcriptomes issued from AD patients's samples issued from IFG (Inferior Frontal Gyrus), STG (Superior Temporal Gyrus ), FP (Frontal Pole) issued from "the Mount Sinai cohort (Wang, Minghui et al. Scientific Data; Sept 2018).

MayoBB_TCX_CPM_ClinicalDiagnosis_AD-NCI

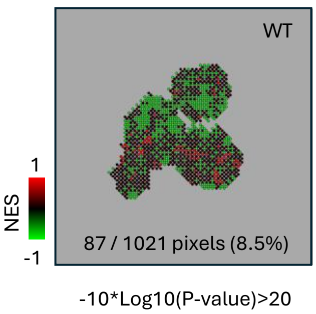


276 subjects : Alzheimer's disease (AD), N=82, progressive supranuclear palsy (PSP), N=84, pathologic aging (PA), N=30, and control (CON), N=80. Allen, Mariet et al; Scientific data vol. 3 160089. 11 Oct. 2016. **Brain region: Temporal Cortex (TCX).** Differential Gene expression analysis from Normalized RNA-seq data: AD vs No cognitive impairment (NCI) control samples.

MayoBB_CBE_CPM_Clinical Diagnosis_AD-NCI

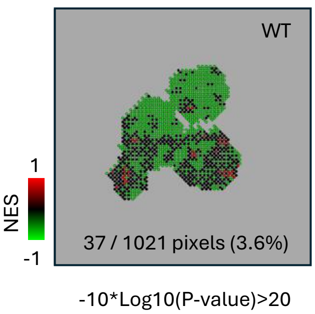


278 subjects: Alzheimer's disease (AD), N=86, progressive supranuclear palsy (PSP), N=84, pathologic aging (PA), N=28, and control (CON), N=80. Allen, Mariet et al; Scientific data vol. 3 160089. 11 Oct. 2016. **Brain region: Cerebellum (CBE).** Differential Gene expression analysis from Normalized RNA-seq data: AD vs No cognitive impairment (NCI) control samples.

MSBB_PHG_CPM_CDR_AD-NCI

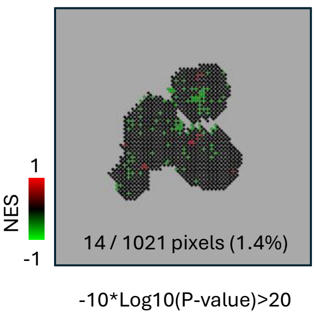


244 subjects: Alzheimer's disease (AD), N=176, No cognitive impairment (NCI), N=68**. Clinical Dementia Rating (CDR): AD vs NCI**. Wang, Minghui et al. "The Mount Sinai cohort of large-scale genomic, transcriptomic and proteomic data in Alzheimer's disease." Scientific data vol. 5 180185. 11 Sep. 2018. **Brain region : Parahippocampal Gyrus (PHG).** Differential Gene expression analysis from Normalized RNA-seq data: AD vs No cognitive impairment (NCI) control samples.

MSBB_PHG_CPM_Braak_B3-B1

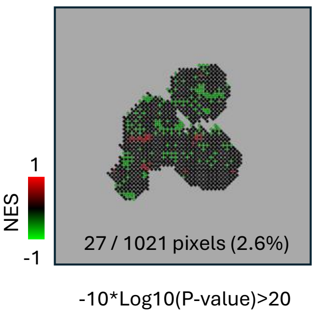


244 subjects: Alzheimer's disease (AD), N=176, No cognitive impairment (NCI), N=68**. Braak Stage** (neuropathological score focused on tau neurofibrillary tangles) B3 (stage V/VI; neocortical) vs B1 (stage 0; normal, I/II (hippocampal) . Wang, Minghui et al. Scientific data vol. 5 180185. 11 Sep. 2018. **Brain region : Parahippocampal Gyrus (PHG).** Differential Gene expression analysis from Normalized RNA-seq data between samples labeled as Braak stage B3 vs B1.

MSBB_PHG_CPM_CERAD_C3-C0

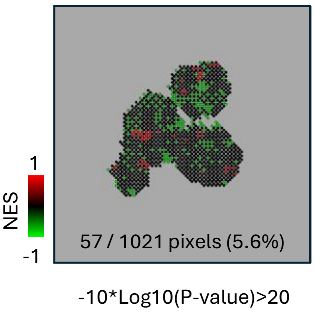


244 subjects: Alzheimer's disease (AD), N=176, No cognitive impairment (NCI), N=68**. CERAD stage** (Neuropathological score focused on the frequency of Beta-amyloid neuritic plaques) C3 (Frequent/Definite AD) vs C0 (None / Not AD). Wang, Minghui et al. "The Mount Sinai cohort of large-scale genomic, transcriptomic and proteomic data in Alzheimer's disease." Scientific data vol. 5 180185. 11 Sep. 2018. **Brain region : Parahippocampal Gyrus (PHG).** Differential Gene expression analysis from Normalized RNA-seq data between samples labelled as CERAD C3 vs C0.

**Supplementary Figure S10. Pixel-level gene set enrichment analysis (GSEA) of AD-upregulated signatures retrieved from Alzheimer DataLENS in WT Brain organoids.** Spatially-resolved transcriptomics (SrT) data assessed from 5-months WT control brain organoids has been interrogated for enrichment of AD-upregulated signatures derived from human AD patient cohorts. These signatures were defined using different disease-stratification strategies, including Clinical Dementia Rating (CDR), Braak stage, and CERAD score, across multiple brain regions and data collections. Each AD-upregulated signature was used as a gene set for pixel-level GSEA of the SrT data. GSEA outcomes were first P-value filtered (-10*Log10(P-value)>20), then expressed as normalized enrichment score (NES) per pixel region (Heatmap). For each of the displayed maps the source of the data samples used as gene set, the analyzed brain region, as well as the AD scoring strategy is described, as retrieved in the DataLENS database.

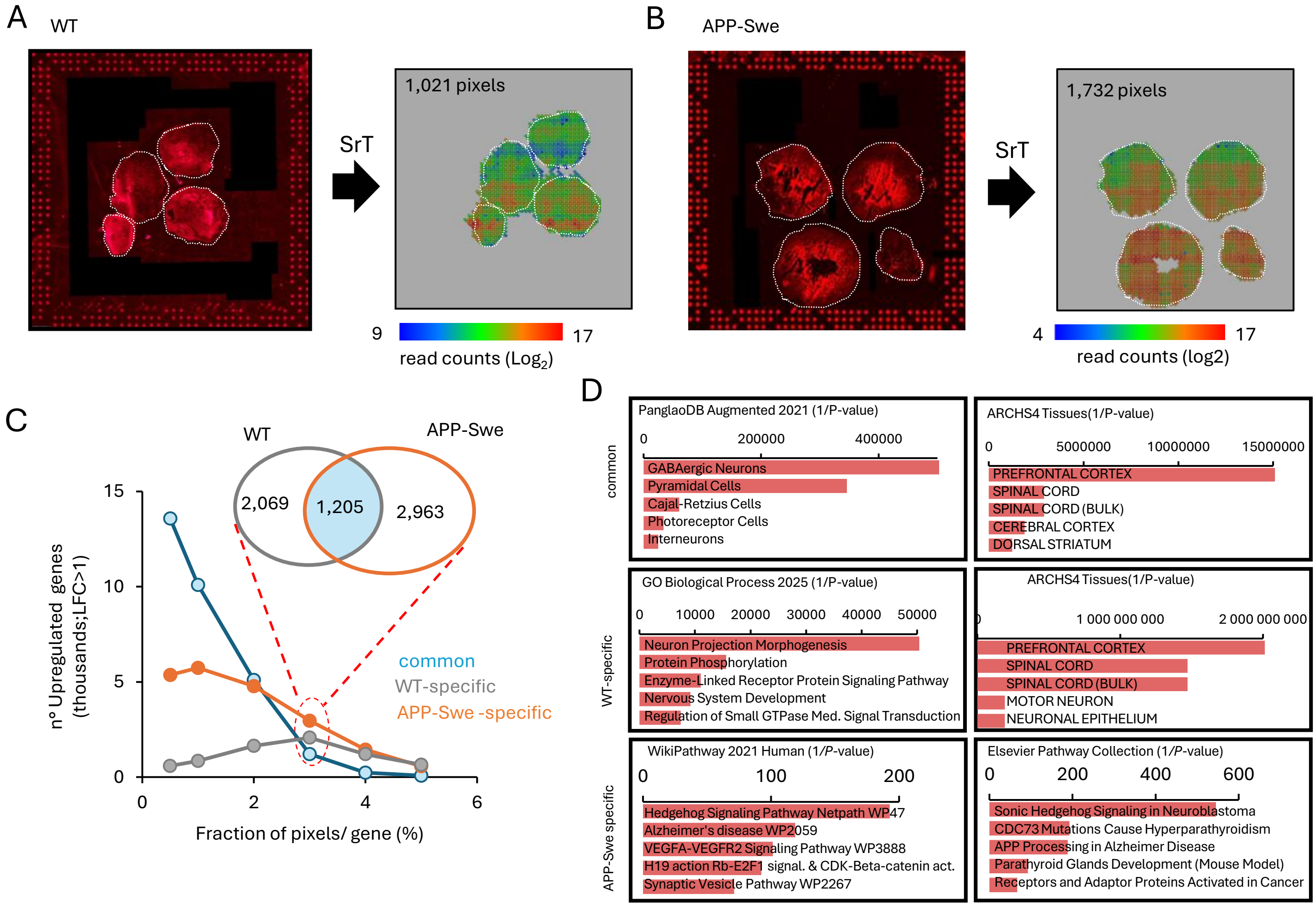


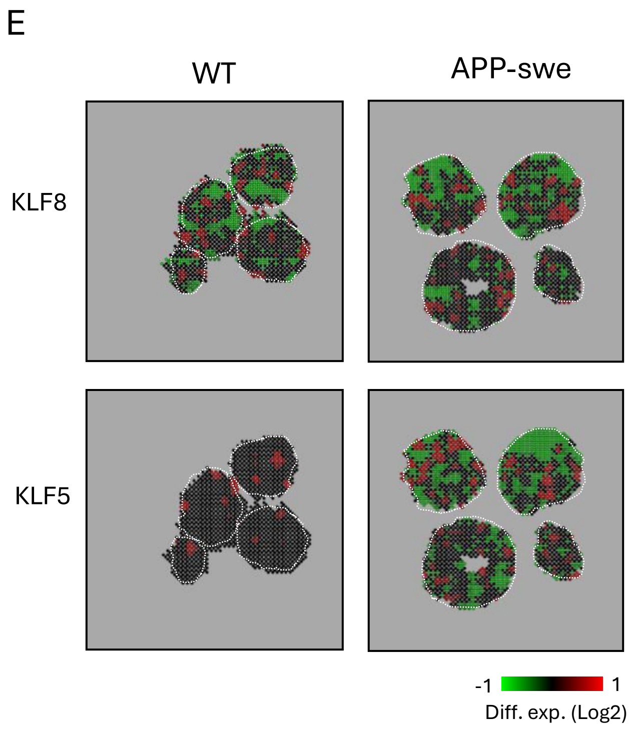


**Supplementary Figure S11. Spatially-resolved transcriptomics (SrT) assay performed on WT and APP-Swedish BORGs of 5 months of age. (A)** Left panel WT: Micrograph displaying cDNA capture in WT BORG sections by in-house manufactured DNA arrays in which the reverse transcriptase assay is performed in presence of dCTP-Cy3. Notice the fiducial borders visualized thanks to the deposition of Cy3-labelled DNA probes. Right panel WT: SrT profile displaying 1,021 pixels corresponding to local transcriptomes. **(B)** Same as (A) but assessed on APP-Swedish BORGs. **(C)** Comparison between the number of up-regulated genes across all pixels in WT and APP-swe SrT data and the fraction of pixels per gene relative to the total number of pixels per profile. Venn-diagram displays the condition where up-regulated pixels per gene correspond to at least 3% of all pixels within the tissue. **(D)** Gene Ontology (GO) analysis performed on the commonly upregulated genes, as well as on the WT or APP-Swedish specific. GO enrichment analysis has been performed with the platform EnrichR (https://maayanlab.cloud/Enrichr/). **(E)** Spatially-resolved differential expression associated to the genes KLF8 and KLF5 in both WT or APP-Swedish tissues. Notice that the differential expression is computed between the read counts per gene assessed per pixel relative to the average read counts over the whole tissue. Despite there are multiple BORGs, they are considered as a single tissue for these analyses.
